# Programmed axon death executor SARM1 is a multi-functional NAD(P)ase with prominent base exchange activity, all regulated by physiological levels of NMN, NAD, NADP and other metabolites

**DOI:** 10.1101/2021.07.14.451805

**Authors:** Carlo Angeletti, Adolfo Amici, Jonathan Gilley, Andrea Loreto, Antonio G. Trapanotto, Christina Antoniou, Michael P. Coleman, Giuseppe Orsomando

## Abstract

SARM1 is an NAD glycohydrolase and TLR adapter with an essential, prodegenerative role in programmed axon death (Wallerian degeneration). It has low basal NADase activity that becomes strongly activated by NAD precursor NMN. Very high levels of NAD oppose this activation, competing for the same allosteric site on SARM1’s regulatory ARM domain. Injury or diseases that deplete axons of NMNAT2, an essential enzyme converting NMN to NAD, cause SARM1 activation. The resulting NAD degradation by SARM1, combined with loss of NAD synthesis by NMNAT2, causes rapid depletion of axonal NAD. This NAD loss is widely assumed to mediate axon death and is consequently a key focus for therapeutic strategies for axonopathies. However, like other NAD(P) glycohydrolases, SARM1 has additional enzyme activities whose contributions, consequences and regulation need to be fully understood. Here, we compare the multiple actions and regulation of recombinant human SARM1 with those of two other NAD(P) glycohydrolases, human CD38 and *Aplysia californica* ADP ribosyl cyclase. We find that SARM1 has the highest transglycosidation (base exchange) activity of these enzymes at neutral pH and with some bases this dominates NAD(P) hydrolysis and cyclisation. Moreover, like its NADase and NADPase reactions, SARM1-mediated base exchange at neutral pH is activated by increases in the NMN:NAD ratio, which we show for the first time can act in the presence of physiological levels of both metabolites. We establish that SARM1 base exchange is the most likely physiological source of calcium mobilizing agent NaADP, and potentially of other NAD(P) analogues, which could contribute to axon and cell death. We also identify regulatory effects of free pyridine bases, of NADP and of nicotinic acid riboside (NaR) on SARM1 that represent further therapeutic opportunities. These data will help to pinpoint which of the multiple functions of SARM1 is responsible for axon degeneration and how it can be optimally targeted to block axon degeneration in disease.

## INTRODUCTION

SARM1 is an intracellular adaptor of Toll-like receptor (TLR) signalling and a prodegenerative enzyme with a central role in programmed axon death, or Wallerian degeneration [1, 2]. Wallerian degeneration occurs when axons are physically transected but it is now clear that the underlying programmed axon death mechanism is also activated by many toxins, gene mutations or metabolic disruption [3, 4]. SARM1 is also constitutively hyperactivated by mutations associated with ALS [5, 6]. The crucial step common to many of these is loss of functional NMNAT2 from axons, an enzyme synthesising the dinucleotide NAD from its precursor mononucleotide NMN. Specific NMNAT2 mutation or knockdown confirms that loss of this single protein is sufficient to cause axon degeneration or axon growth failure that is absolutely dependent on SARM1, with complete rescue when SARM1 is removed [7-10]. Thus, loss of NMNAT2 lies upstream of the role of SARM1 in programmed axon death. Surprisingly, the rise in NMNAT2 substrate, NMN, when this enzyme is depleted plays a particularly important role in activating the pathway [11-14]. The unexpected discovery that SARM1 has intrinsic NADase activity [15] then led to the finding that NMN activates SARM1 NADase [16]. NMN is now known to do this by binding an allosteric site on the SARM1 inhibitory ARM domain, and high levels of NAD compete to bind the same site, thereby opposing activation by NMN [17-19]. VMN, a metabolite of the neurotoxin and disused rodenticide vacor, binds the same site and activates SARM1 even more potently [20]. The current working model of the programmed axon death signalling pathway is shown in Figure 1A.

**Figure 1.**
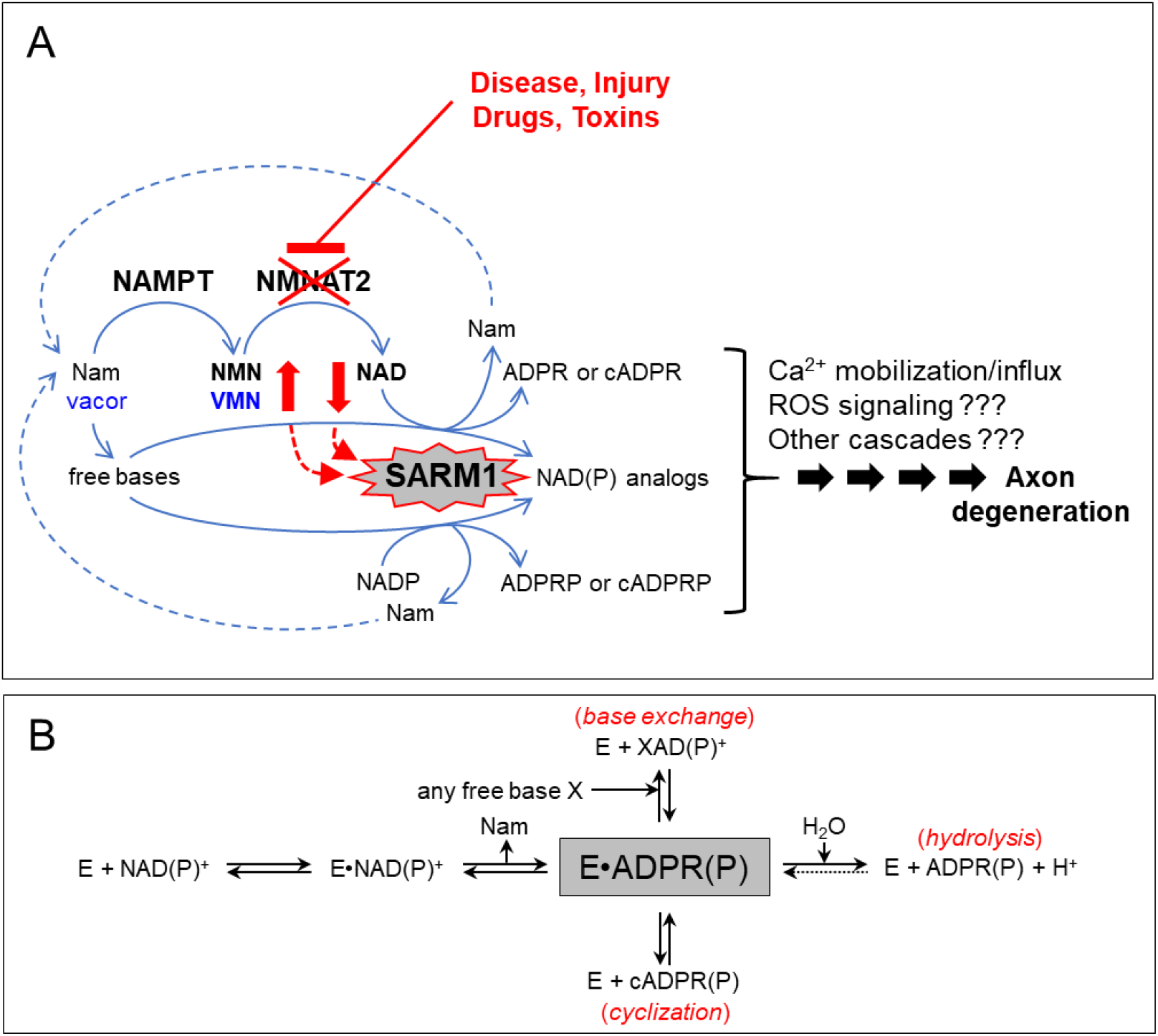
SARM1 is a central executioner of axon death via a multicomposite catabolic reaction typical of multifunctional NAD(P) glycohydrolases. (**A**) Programmed axon degeneration is a widespread axon death mechanism driven by activation of SARM1 NAD(P)ase and prevented by its negative regulator NMNAT2, a short half-life enzyme essential to convert NMN into NAD. NMNAT2 forms together with the upstream enzyme NAMPT a key two-step pathway for NAD salvage in mammals that also provides metabolic conversion of the prodrug vacor into the neurotoxic, recently discovered, VMN intermediate. When NMNAT2 in axons becomes depleted or inactive, both NMN and VMN rise and NAD declines concomitantly; these fluctuations trigger SARM1 NAD(P)ase and, as shown here also base exchange, and thus initiate a signaling cascade that culminates into axon death. The early activation mechanism and key components of this process are highlighted in red and in blue for vacor toxicity. (**B**) Ordered Uni-Bi reaction mechanism of multifunctional NAD(P)ases (EC 3.2.2.6) like SARM1. From the substrate NAD(P) the pyridine moiety, Nam, is released first and then two distinct ADP ribosylated products arising from a single common intermediate (grey boxed). These may be ADPR(P) via classical hydrolysis or cADPR(P) via anhydrous cyclization. A third product XAD(P) represents any dinucleotides formed instead via transglycosidation (EC 2.4.99.20), a reaction of base exchange that replaces the pyridine moiety of the substrate with any related free base available.

Another NAD(P) glycohydrolase, CD38, shows some catalytic similarity to SARM1 but no sequence similarity [3, 15]. CD38 NAD(P)ase does not appear to influence axonal degeneration [21] although it has roles in intracellular NAD homeostasis and calcium signaling [22] and is linked to other human diseases [23]. SARM1 and CD38 thus control distinct fates in mammalian cells via intervention on related metabolism [24], suggesting there may be subtle but important differences in the specific reactions they catalyse, categorised as NAD hydrolysis (EC 3.2.2.5), multifunctional enzymes catalysing both synthesis and hydrolysis of cADPR (EC 3.2.2.6), or NADP hydrolysis and base exchange (EC 2.4.99.20) (Fig. 1B). The structural basis of these distinct activities is at least partially understood [25]; it involves an active site glutamate residue [15, 26, 27], which transiently accommodates the ADPR moiety after hydrolysis of the β-N-glycosidic bond and release of nicotinamide (Nam). The energy from this bond is conserved within the enzyme-bound complex allowing the direct formation of multiple products depending on which moiety is available or effectively joins the catalytic pocket. Use of water leads to hydrolysis, free pyridines lead to transglycosidation (base exchange) and in the absence of both there is cyclisation [26]. All of these activities have important roles in the synthesis of calcium mobilisers [24, 28]: ADPR is the product of NAD hydrolysis, cADPR of its cyclisation and base exchange is pivotal for the synthesis of NaADP from nicotinate and NADP (EC 2.4.99.20), of which there is no other known source in mammals [29].

In healthy axons, SARM1 exhibits a low, basal NAD(P)ase activity [28]. Removing SARM1 greatly lowers basal levels of one NAD-derived product, cADPR, in several neural tissues but has only a modest, if any, effect on NAD levels [9, 28, 30]. Conversely, basal NAD is strongly elevated by removal of CD38 [21], but with limited effect on cADPR [31]. This suggests that while CD38 is responsible for most NAD degradation under basal conditions, its primary product *in vivo* is not cADPR, while SARM1 is the main regulator of basal cADPR but other, possibly greater activities of SARM1 remain possible. Base exchange by SARM1 has been reported very recently in live cells [32] but, other than it being unaltered by NMN at pH 4.5 [16], little is known about this activity. The complex, tripartite domain structure of SARM1 contrasts with CD38 and its homologue ADP ribosyl cyclase from *Aplysia californica* (*Aplysia* cyclase) which lack separate regulatory domains [33, 34]. SARM1 has an N terminal ARM domain for auto-inhibition, two internal SAM domains for multimerization, and a C-terminal TIR domain for catalysis [18, 19]. TIR is an evolutionary ancient domain with many roles in innate immunity via protein interactions [35], which appears to have independently evolved as a NAD(P)ase [36]. Hence, SARM1’s catalysis appears to be a product of functionally convergent evolution with homodimeric *Aplysia* cyclase and CD38 but within a multidomain protein architecture making it ideally suited as a proposed molecular switch [37] from NAD(P) homeostasis in healthy axons to self-reinforcing NAD(P) decline in axons damaged beyond repair.

Despite the well-supported working model of programmed axon death (Fig. 1A), many questions remain. One concerns the direct cause of axon death downstream of SARM1 activation. NAD depletion, leading to loss of ATP synthesis and of many other NAD-dependent activities [38], is an attractive candidate mechanism that has been widely assumed to be correct but direct evidence for this is lacking. Neurons and their axons can survive with remarkably low NAD levels [17], and while one SARM1 product, cADPR, has been partially excluded as a direct cause of axon death [28], there are others (see Fig. 1A) that have not been investigated [16]. There are also other pyridine nucleotides degraded by SARM1 [36]. Moreover, while the NMN:NAD ratio is able to influence NAD turnover and cADPR synthesis by SARM1, its effect on these other activities especially at neutral pH remains unknown. It also remains unclear how SARM1 responds to modulation of either of these two key regulators when both are present at physiologically-relevant levels, and whether other NAD-related metabolites also influence activity.

With all this in mind, we have compared the enzyme activities and regulation of hSARM1 to those of hCD38 and *Aplysia* cyclase *in vitro*. First, we identify a strong bias of SARM1 towards base exchange, strongly suggesting this newly identified, multifunctional NAD(P)ase as a potential key regulator *in vivo* of the potent Ca^2+^ mobilizer NaADP. Second, we show the ratio of NMN:NAD is a key determinant of SARM1 activity at physiologically-relevant levels of each metabolite, an effect that requires the ARM domain and that at neutral pH is true for all known enzyme activities of SARM1. Third, we show there is also regulation by NADP through TIR domain interaction, again at physiologically-relevant concentrations. Finally, we identify selective active site inhibition by nicotinic acid riboside (NaR) and analogues that could underlie new therapeutic strategies for axonopathies.

## RESULTS

### Preliminary *in vitro* assays on selected multifunctional NAD(P)ases

We first established appropriate incubation times to determine initial rates with saturating NAD for commercial preparations of human CD38 (hCD38) and *A. californica* ADP ribosyl cyclase (*Aplysia* cyclase), and of human SARM1 full length (hSARM1) isolated from HEK cells (Fig. S1), assaying the generation of multiple products by HPLC. We established specific activities for basal NADase through hydrolysis or cyclisation combined of ∼0.02 U/mg for hSARM1, ∼7 U/mg for hCD38, and ∼50 U/mg for *Aplysia* cyclase. Given the absence of free pyridines we could individually measure the hydrolysis and cyclization activity of the ordered Uni-Bi mechanism [39] through the release of free or cyclic ADP ribosyl moiety (Fig. 1B). There was clear divergence between the products of the three enzymes (Fig. 2A), indicating a strong preference for cyclization by *Aplysia* cyclase and hydrolysis for hCD38 accounting for >98% of NAD consumption in both cases. hSARM1 showed more mixed activity with hydrolysis accounting for ∼90% NADase activity and cyclization ∼10% (Fig. 2A). Varying NAD concentrations had no effect on these proportions (not shown).

**Figure 2.**
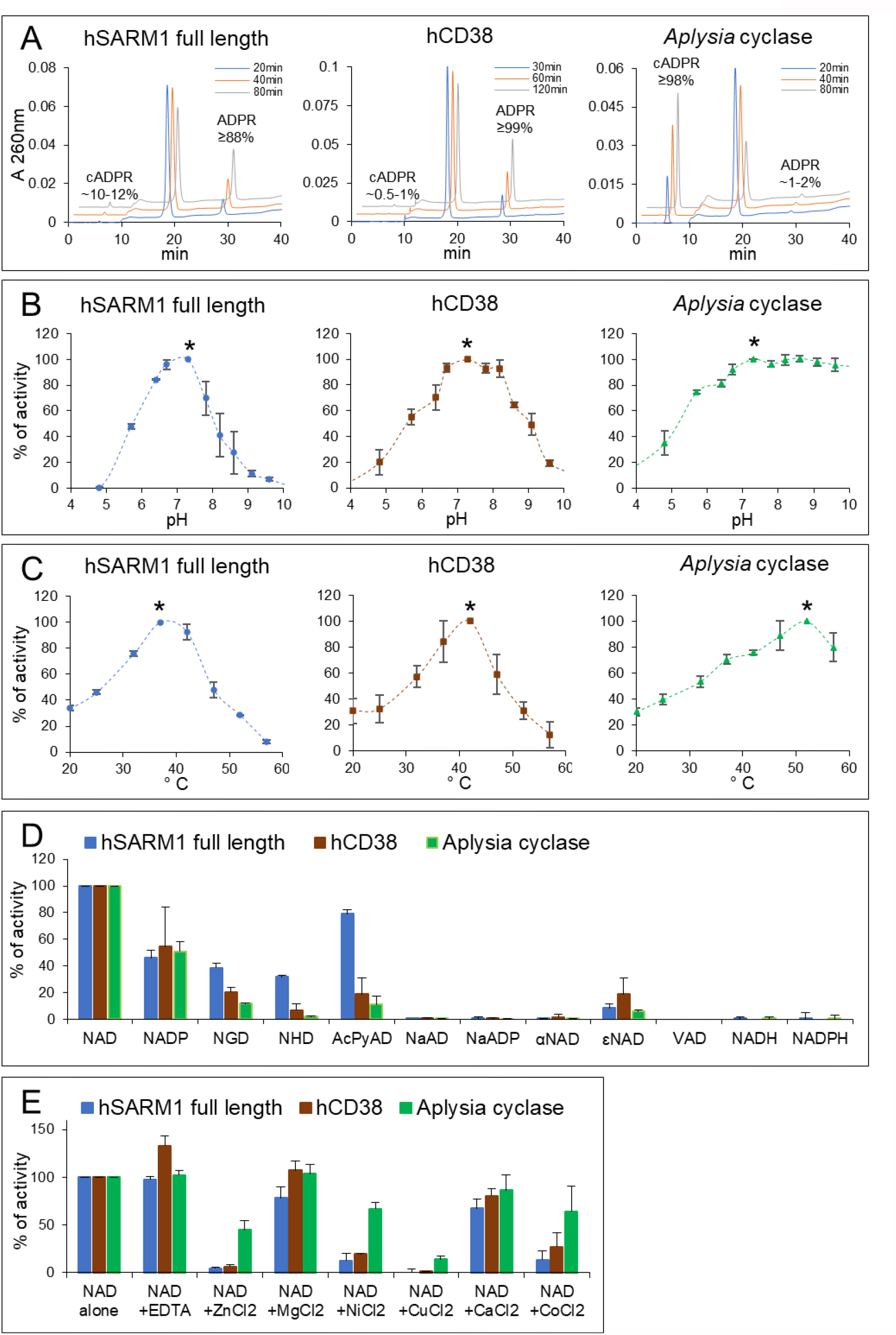
Preliminary characterization of selected multifunctional NAD(P)ases. (**A**) C18-HPLC UV profiles from time-course analyses of the three indicated multifunctional NAD(P)ase enzymes studied, all assayed with 250 µM NAD at 25 °C under similar rates of substrate consumption. Both ADPR and cADPR products accumulating into diverse proportions are highlighted while unmarked in between, the substrate NAD peak declines in parallel and proportionally by time. (**B**) pH studies carried out by HPLC assays in universal buffer (Tris/Bis-Tris/Na-acetate) for 1 hour at 25 °C using 0.25 mM NAD and 7 µg/ml hSARM1 or 0.5 mM NAD and 0.07 µg/ml CD38 or 6 mM NAD and 0.08 µg/ml *Aplysia* cyclase. Data are Mean ± SEM from n = 3 and are normalized to relative maxima of each curve (see asterisks). (**C**) Optimum temperatures. HPLC assays were set for 1 hour using 0.25 mM NAD and 11 µg/ml hSARM1 or 0.5 mM NAD and 0.1 µg/ml CD38 or 2.5 mM NAD and 0.14 µg/ml *Aplysia* cyclase. Data are Mean ± SEM from n = 3 and are normalized as in B. (**D**) Preferred substrates and (**E**) effects of metal ions. Various dinucleotides (250 µM each) or metal ions (1 mM each) were assayed at 25 °C by HPLC for 2-6 hours using 5.5 µg/ml hSARM1 or 0.1 µg/ml CD38 or 0.028 µg/ml *Aplysia* cyclase. Data are Mean ± SEM from n ≥ 2. Dinucleotide analogs indicated are: NGD, nicotinamide guanine dinucleotide; NHD, nicotinamide hypoxanthine dinucleotide; AcPyAD, 3-acetylpyridine adenine dinuclotide; NaAD, nicotinic acid adenine dinuclotide; NaADP, NaAD phosphate; αNAD, alpha-NAD; εNAD, nicotinamide 1,N^6^-etheno adenine dinucleotide; VAD, vacor adenine dinucleotide; NADH and NADPH, reduced NAD and NADP.

We determined pH and temperature optima for each enzyme under comparable conditions. Maxima for hSARM1 and hCD38 were similar, within the physiological range of mammals (Fig. 2B,C), whereas *Aplysia* cyclase was more tolerant of other conditions, probably reflecting its marine invertebrate origin, remaining active at basic pH or high temperatures (Fig. 2B,C). At pH 5 and below, where CD38 shows base exchange activity with a possible physiological role in lysosomes [24, 40], both hCD38 and hSARM1 showed NADase activities of ∼20% or less of their maxima (Fig. 2B). All three enzymes at 25 °C showed activity of ∼40% of their maxima (Fig. 2C), so consistent with other recent reports [15, 16, 36] this temperature and pH 7.5 were chosen for subsequent experiments.

We then tested the use of alternative substrates and effects of divalent metal cations (Fig. 2D,E) in view of the reported influence of zinc ions on SARM1 activity [41] and also to determine assay conditions suitable to discriminate hSARM1 and hCD38 activity in mammalian tissue extracts, analogous to our reported isoform-specific assays for NMNAT activities [42]. The physiological redox dinucleotides NAD and NADP were both among the preferred substrates of all three enzymes based on consumption rates. In contrast, there was negligible usage of either the corresponding reduced forms, NADH and NADPH, or the deamidated forms, NaAD and NaADP (Fig. 2D). There was some consumption of NGD and NHD, which also occur naturally [43], and of AcPyrAD and the artificial substrate εNAD. However, there was no activity towards αNAD or the vacor derivative VAD, despite prolonged incubations. Regarding the metal salts, magnesium, or chloride counterions had no effect, while most other divalent cations tested, particularly copper and zinc, showed clear inhibition (Fig. 2E). These results are consistent with previous reports [36, 41, 44, 45], but this side-by-side comparison reveals broader substrate use and greater divalent cation inhibition for hSARM1 compared to hCD38. The relative rates of hydrolysis and cyclisation were also broadly similar to those with NAD (Fig. 2A), except that hSARM1 showed a lower cyclisation rate for NADP (≤1% of total NADP consumption) and CD38 a much higher cyclisation rate for NGD (∼80% of total NGD consumption) [27, 46]. These features, together with a selective inhibition by pyridine ribosides as described below, could be the basis of assays to distinguish CD38 and SARM1 in complex mixtures.

### Kinetics studies towards structural regulations on multidomain hSARM1

We then studied kinetics of these three enzymes with respect to their physiological substrates NAD and NADP. For NAD, both hCD38 and *Aplysia* cyclase showed classically hyperbolic curves (Fig. 3A) as reported [39, 47], whereas hSARM1 did not. Its NADase activity rose to a maximum at 0.25 mM NAD and then progressively fell to almost zero at 4 mM (Fig. 3A left). Measurements of NAD extracted from most mammalian tissues indicate levels between 0.2-1.0 mM [48] so the optimum is well positioned to respond to intracellular fluctuations of NAD such as those in circadian rhythms [49]. This is particularly so in brain where basal levels are estimated at 0.3 mM [48]. The absence of such strong inhibition on N-terminally-truncated SARM1, consisting of only SAM and TIR domains (Fig. 3B and Table 1) is consistent with recent reports of allosteric binding of NAD to the regulatory ARM domain [17-19], publications which reported widely-varying inhibitory effects of NAD alone, or of blocking of NMN-induced activation by NAD. Importantly, our experiments used preparatory FPLC to guarantee the absence of contaminating impurities that often arise by degradation of NAD, including NMN and Nam, which will interfere with the measured response to variable NAD (see below).

**Figure 3.**
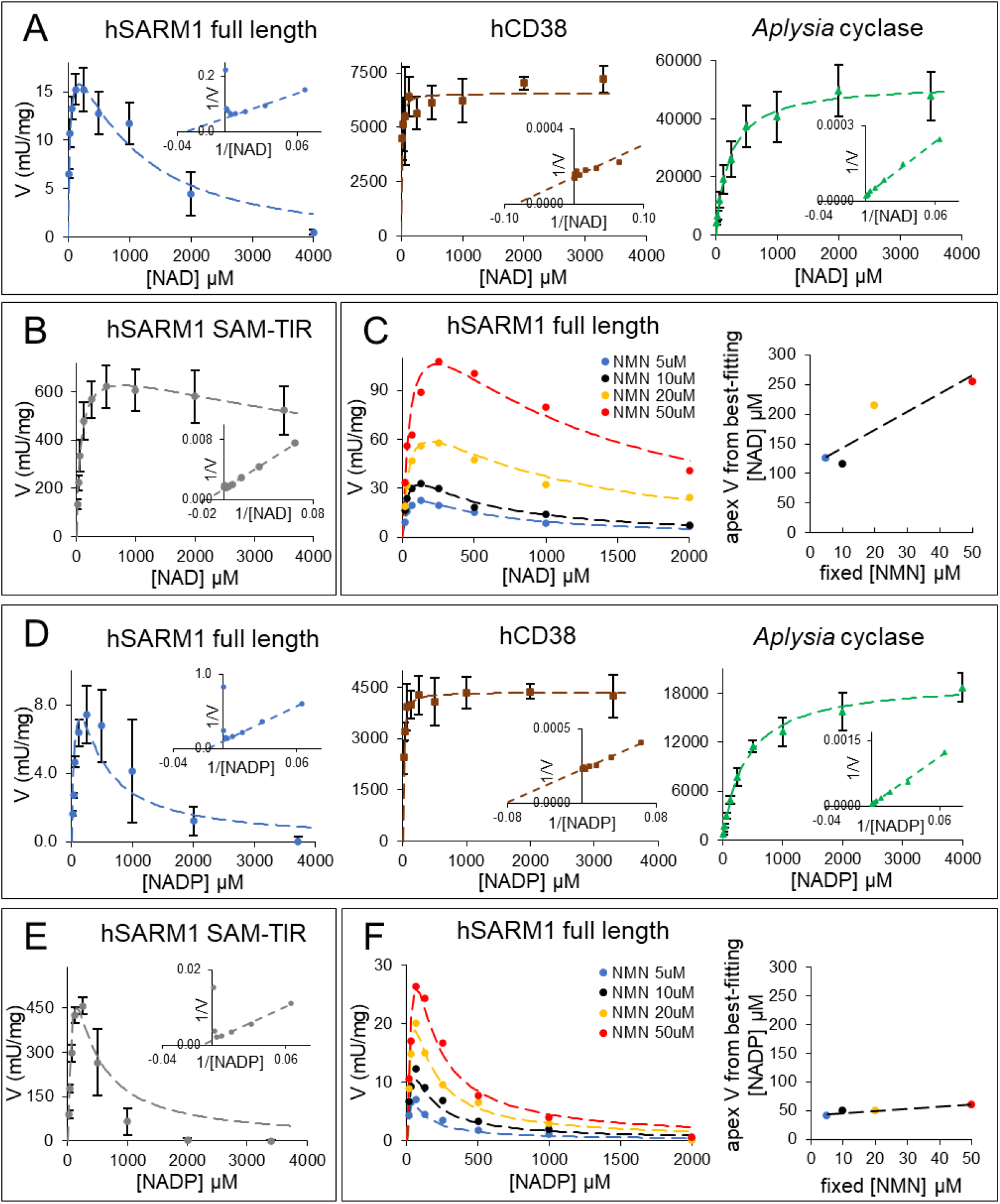
NAD and NADP kinetics by SARM1 in comparison to other multifunctional NAD(P)ases and under allosteric triggering by NMN. (**A**) NAD kinetics of the three multifunctional NAD(P)ases. Initial rates were measured by HPLC using 3 µg/ml hSARM1 or 0.05 µg/ml CD38 or 0.02 µg/ml *Aplysia* cyclase. Assays were carried out at 25 °C for 30 min. Data are Mean ± SEM from n ≥ 3. Insets, double reciprocal plot analyses. Dotted lines, best fitting analyses carried out using equations 1 and 3 in Methods. The recalculated kinetic parameters are shown in Table 1. (**B**) NAD kinetics of the human SARM1 fragment SAM-TIR with constitutive NAD(P)ase that is not inducible by NMN because of the lack of the auto-inhibitory N-terminal regulatory domain ARM. HPLC assays were carried out at 25 °C for 1 hour using 2.75 µg/ml enzyme per mix. Data are Mean ± SEM from n = 4. Inset, double reciprocal plot analysis. Dotted line, best fitting results from equation 2 in Methods (see also recalculated parameters in Table 1). (**C**) NAD kinetics of hSARM1 full length (3 µg/ml per mix as in A) at increasing micromolar concentrations of the allosteric regulator NMN. Best fitting analysis was carried out on individual curves using equation 3 in Methods to calculate the kinetic parameters shown in Table 1, and the relative curve maxima that were subsequently re-plotted on the flanking graph. Their linear relationship with the trigger amount in each curve indicates competition between NMN and NAD for opposing regulation of hSARM1 NADase. (**D**) NADP kinetics presented as in A above but done by assaying 14.6 µg/ml of hSARM1 or 0.05 µg/ml of CD38 or 0.07 µg/ml of *Aplysia* cyclase at 25 °C for 60-120 min. Data are Mean ± SEM from n = 3. (**E**) NADP kinetics as in B above but done by assaying 1.9 µg/ml of SAM-TIR at 25 °C for 1 hour. Data are Mean ± SEM from n = 2. (**F**) NADP kinetics as in C above (3.7 µg/ml hSARM1 full length per mix) at increasing micromolar concentrations of NMN. Flanking graph, best fitting maxima re-plotted showing NADP effects unrelated to allosteric triggering by NMN of hSARM1 NADPase.

**Table 1.** Kinetic parameters from best fitting analysis of the initial rates shown in Figure 3.

|  |  | NAD substrate |  |  |  |  | NADP substrate |  |  |  |
| --- | --- | --- | --- | --- | --- | --- | --- | --- | --- | --- |
|  |  | catalytic site |  |  | allosteric site |  | catalytic site |  |  | allosteric site |
| | rep (n) | $K_m$ (μM) | $V_{max}$ (mU/mg) | $K_{is}$ (μM) | $K_{ia}$ (μM) | rep (n) | $K_m$ (μM) | $V_{max}$ (mU/mg) | $K_{is}$ (μM) | $K_{ia}$ (μM) |
| hCD38 | 3 | 10.4 ± 2.90 | 6649 ± 124.7 | - | - | 3 | 11.7 ± 0.88 | 4376 ± 24.1 | - | - |
| *Aplysia* cyclase | 3 | 212 ± 13.7 | 51353 ± 1237 | - | - | 3 | 384 ± 0.17 | 20043 ± 724 | - | - |
| hSARM1 SAM-TIR | 4 | 69.5 ± 2.31 | 723 ± 13.7 | >8000 | - | 2 | 83.5 ± 11.9 | 582 ± 46.9 | 151 ± 34.1 | - |
| hSARM1 full length | 4 | 30.3 ± 5.18 | 22.4 ± 5.24 | >8000 | 324 ± 24.6 | 3 | 66.9 ± 32.6 | 14.3 ± 3.14 | 132 ± 49.5 | >8000 |
| +NMN 5μM | 1 | 44.1 | 39.7 | >8000 | 385.1 | 1 | 29.3 | 15.7 | 52.7 | >8000 |
| +NMN 10μM | 1 | 34.2 | 54.1 | >8000 | 414.9 | 1 | 45.9 | 29.9 | 57.4 | >8000 |
| +NMN 20μM | 1 | 44.6 | 88.2 | >8000 | 787.0 | 1 | 45.0 | 52.2 | 60.0 | >8000 |
| +NMN 50μM | 1 | 56.1 | 161 | >8000 | 1201.7 | 1 | 91.7 | 98.1 | 48.0 | >8000 |

Best fitting was achieved by the equations described in the Methods that were applied on replicates as indicated or on single curves at distinct NMN concentrations. “ $K_{is}$ ” and “ $K_{ia}$ ” are the affinity constants for the inhibition exerted, respectively, on the catalytic site on TIR domain and on the allosteric site on ARM domain of hSARM1.

As NMN and NAD exert reciprocal positive and negative regulation of SARM1 through competition at the same allosteric site [17], it is essential to determine SARM1 NADase kinetics close to the physiological levels of each of these regulators, and so far this has not been reported. Thus, we repeated our above analysis of NAD kinetics on full length hSARM1 in the presence of increasing amounts of NMN (Fig. 3C). In particular, we sought to determine the degree to which physiologically-relevant levels of NAD could counter activation by physiological NMN and how much falling NAD contributes to SARM1 activation around these physiological levels. We find that while inhibition by very high, non-physiological levels of NAD remains when hSARM1 is activated by NMN, NAD concentrations close to those in brain had only a marginal effect in countering activation by NMN as it rises from its physiological level of ca. 6 µM in brain [48] to more than double that in compromised axons [14] (Fig. 3C). In lesioned nerves, NAD levels approximately halve before axon fragmentation occurs [14] although it is possible that a larger decline in axons is partially masked by relatively stable NAD in glia. Figure 3C shows that halving NAD from its normal nervous system level of ca. 300 µM marginally increases hSARM1 activity when NMN is close to basal levels (5 µM) but has decreasing effect as NMN rises. Indeed, if axonal NAD does drop by more than 50% this could even lower hSARM1 NADase due to the lower availability of substrate. The shift in curve maxima to the right (Fig. 3C) and increasing *K*_ia_ (Table 1) with increasing NMN are consistent with competition by NMN and NAD for the same regulatory site as reported [17-19, 37]. These data strongly suggest that the rise in NMN when NMNAT2 is compromised is the principal determinant of hSARM1 activation, while fluctuations in NAD within the physiological or pathological range have limited effect.

SARM1 also has NADPase activity [36]. Considering the role of its reduced form in ROS buffering as well as many anabolic reactions, NADP loss is an additional candidate mechanism for SARM1-dependent axon death, but the response of SARM1 NADPase to NMN, or indeed to NADP itself, has not been reported. Thus, we performed similar analysis to those described above for NAD but now with variable NADP and NMN (Fig. 3D-F and Table 1). Again we saw substrate inhibition for full length hSARM1 NADPase (Fig. 3D left), but in contrast to our findings with NAD this effect was fully retained in the absence of the ARM domain (Fig. 3B,E). Physiologically-relevant levels of NMN also strongly activated SARM1 NADPase (Fig. 3F) with a potency very similar to that for SARM1 NADase (Fig. S2) and the absence of a rightwards shift of the NADPase maxima, or of increasing *K*_is_ as NMN rises, further indicate that inhibition by NADP is not mediated by competition for the NMN binding site of the ARM domain. Instead, it is likely to interfere locally with TIR catalysis. Importantly, substrate inhibition by NADP shows a kinetic constant (*K*_is_) of just around 50 µM, which again is close to best estimates of physiological NADP concentration in most mammalian cells [50] and tissues like brain (unpublished personal data).

In summary, these data show that (1) both NAD and NADP inhibit SARM1 TIR catalysis at high concentrations; (2) NADP is the more potent inhibitor (*K*_is_ ∼50 uM vs *K*_ia_ ∼300 uM), making this physiologically relevant even at its lower *in vivo* concentration; (3) inhibition by NAD is principally allosteric, in competition with NMN, whereas that by NADP is likely to occur at the catalytic site and is independent of NMN.

### Dominant and unique base exchange capability of hSARM1 NAD(P)ase

The base exchange activity of SARM1, using NADP and Na as substrates, was previously shown to exceed that of NAD hydrolysis or cyclisation [16]. However, it was not further activated by NMN under the acidic assay conditions used to mirror the proposed NaADP synthesis activity of CD38 in lysosomes [40]. SARM1 is not thought to be a lysosomal enzyme [24] so it is important to extend these studies to neutral pH and to other bases, and to directly compare base exchange and hydrolysis/cyclisation rates using the same dinucleotide substrate. Thus, we characterised SARM1 base exchange activity for NAD or NADP paired with three different pyridine bases: 3-acetyl pyridine (AcPyr), vacor (1-(4-nitrophenyl)-3-(pyridin-3-ylmethyl)urea), and nicotinic acid (Na) at pH 7.5 (Fig. 4, Table S1). Both hCD38 and *Aplysia* cyclase showed much lower base exchange reactions than hSARM1 under these conditions, and we confirmed previous reports about their undetectable activity at this pH with Na in particular (Fig. 4A left, middle panels). In contrast, for both full length SARM1 and its ARM-lacking fragment (Fig. 4A right, 4B left), base exchange was detectable for all three bases, including Na, and we also found that (1) the base exchange activity can displace hydrolysis and cyclisation for both NAD and NADP to a degree dependent on the base used, and (2) overall use of both NAD and NADP increases substantially in the presence of bases when the ARM domain is present. The presence of AcPyr almost abolishes other reactions of SARM1 while vacor and Na leave both NADase and NADPase activities little altered although some extra VAD(P) or NaAD(P) are formed from exchanges, respectively accounting for 20-30% or 2-3% of total products (Fig. 4B left, Fig. S3, Table S1). All of these activities increased further still in the presence of either NMN or VMN [20] and leveled off at a plateau of ∼8-10 fold (Fig. 4B middle, right panels). Thus, base exchange at neutral pH is regulated similarly to NAD(P) hydrolysis and cyclisation but that activation by these regulators is not additive with the enhanced NAD or NADP loss caused by the free base alone. Lastly, we confirmed that all three bases above may also work together in mixture at micromolar concentrations (Fig. S4), thus showing how multiple base exchanges may occur for SARM1 at the same time and without any apparent mutual competition among bases.

**Figure 4.**
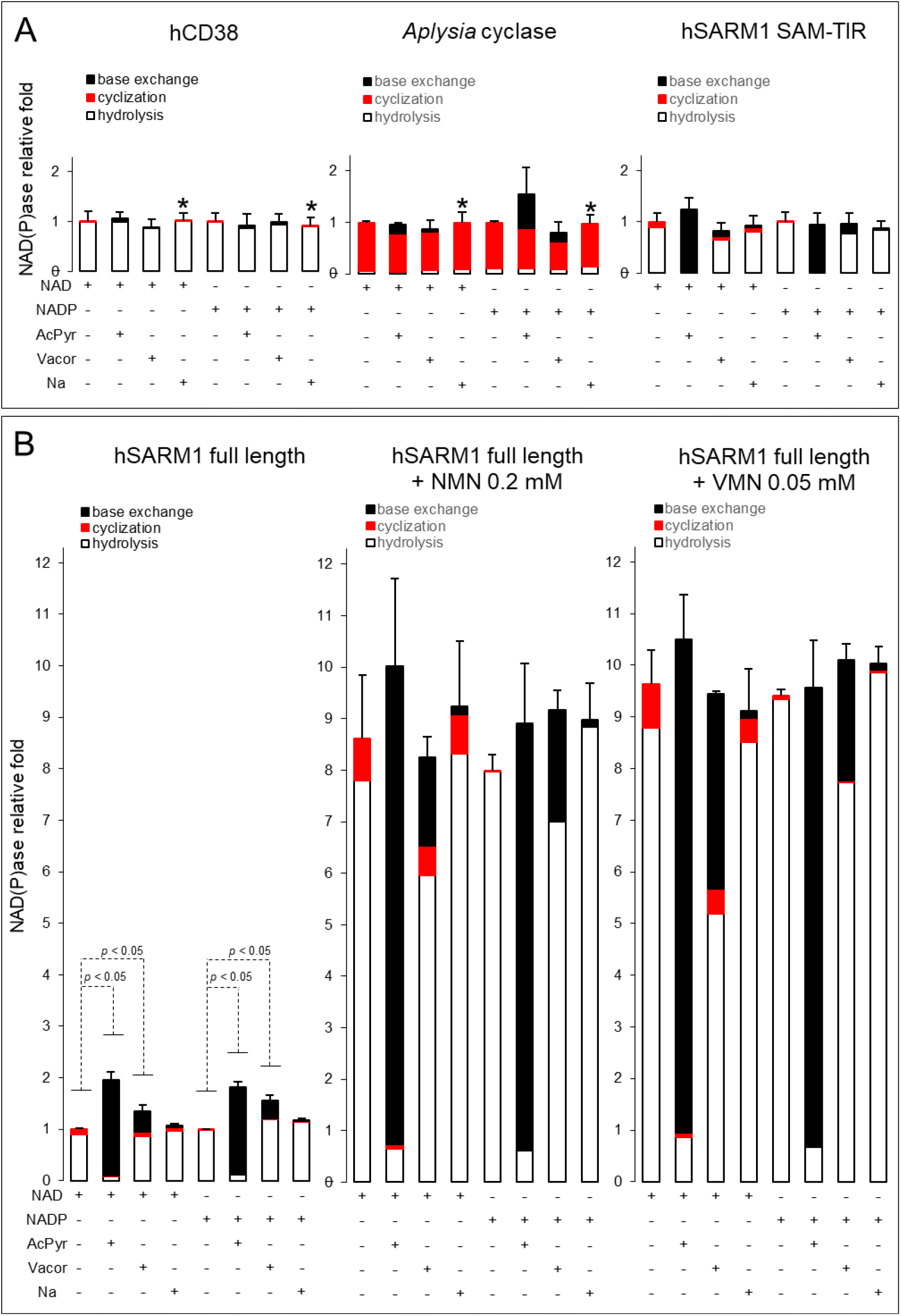
Base exchange reactions typically catalysed by SARM1 at neutral pH. Comparison of base exchange reactions catalyzed in 50 mM HEPES/NaOH pH 7.5 by the indicated multifunctional NAD(P)ases. Rates were measured at 25 °C by HPLC using 0.05 µg/ml CD38 or 0.1 µg/ml *Aplysia* cyclase or 0.5 µg/ml SARM1 SAM-TIR (**upper panel A**) or 7 µg/ml SARM1 full length (**bottom panel B**). Both NAD and NADP substrates were fixed at 250 µM. The free bases 3-acetyl pyridine (AcPyr) and nicotinic acid (Na) were added at 2 mM final. Vacor was added at 0.5 mM given its low solubility at physiological pH. The assay in B (left panel) is also shown in Supplementary (see Fig. S3 with corresponding C18-HPLC UV profiles) and was replicated in the presence of two known allosteric regulators of SARM1, NMN 0.2 mM and VMN 0.05 mM (**B, middle and right panels**), leading both to a maximum triggering effect on SARM1 activity at this concentration as reported [20]. Multiple time stops from individual assays as above were analyzed for linearity, then extents of hydrolysis (white bars), cyclization (red bars) or base exchange (black bars) were calculated from each corresponding product as shown in Fig. 1B and Methods. In detail, base exchanges led to form AcPyrAD from AcPyr, VAD from vacor, NaAD from Na in the presence of NAD or AcPyrADP from AcPyr, VADP from vacor, NaADP from Na in the presence of NADP (see Fig. S3). Rates (Mean ± SEM, n = 2) are shown in histograms for comparison, referred to either NAD or NADP alone controls (arbitrarily fixed to 1). The whole data set is also shown in Table S1. Asterisks (*), indicate conditions where base exchange was below detection.

These data show that base exchange is largely unique to SARM1 in a physiological setting and can be dominant over the hydrolysis or cyclisation of both NAD and NADP. This suggests important physiological roles of SARM1 NAD(P)ase that have so far been little considered and indicates promising new therapeutic strategies (see Discussion).

### Selective inhibition by pyridine ribosides

The NAD(P)ase-enhancing effect on full length hSARM1 exerted by free pyridine bases, and the opposing effects exerted by pyridine mono- and dinucleotides, prompted us to investigate also the effects of other pyridine moieties on this enzyme. So we assessed the pyridine ribosides, also in view of their membrane permeability that could support use as experimental tools and in drug development. We initially tested several natural or artificial pyridine ribosides at fixed concentration on the NADase activity of all three enzymes. To our surprise, we found that two vitamin B3 precursors, NaR and NR [51-53], exerted inhibitory effects but with opposing selectivity for hSARM1 *vs* both hCD38 and *Aplysia* cyclase (Fig. 5A). Vacor derivative VR also strongly inhibited hSARM1 similar to NaR (Fig. 5A). Preliminary IC50 measured at saturating NAD for NaR and VR on full length hSARM1 were 87 and 154 µM respectively, and inhibition of hSARM1 lacking its ARM domain was slightly stronger at IC50s of 36 and 61 µM (Fig. 5B), indicating that this is not allosteric regulation through the ARM domain. Then, using the hSARM1 SAM-TIR fragment, we sought to determine the dissociation constant describing the binding affinity between the inhibitor and the enzyme and the kinetic mechanism of inhibition with results in the low micromolar range (*K*_i_ of 15 µM for NaR or of 25.9 µM for VR) and mixed-type inhibition for both NaR and VR (Fig. 5C,D). We also obtained non-linear secondary plots of slopes and intercepts against concentrations of both NaR and VR, indicating concomitant binding at multiple sites for these inhibitors (Fig. 5C,D right panels with the indicated “n” values > 1).

**Figure 5.**
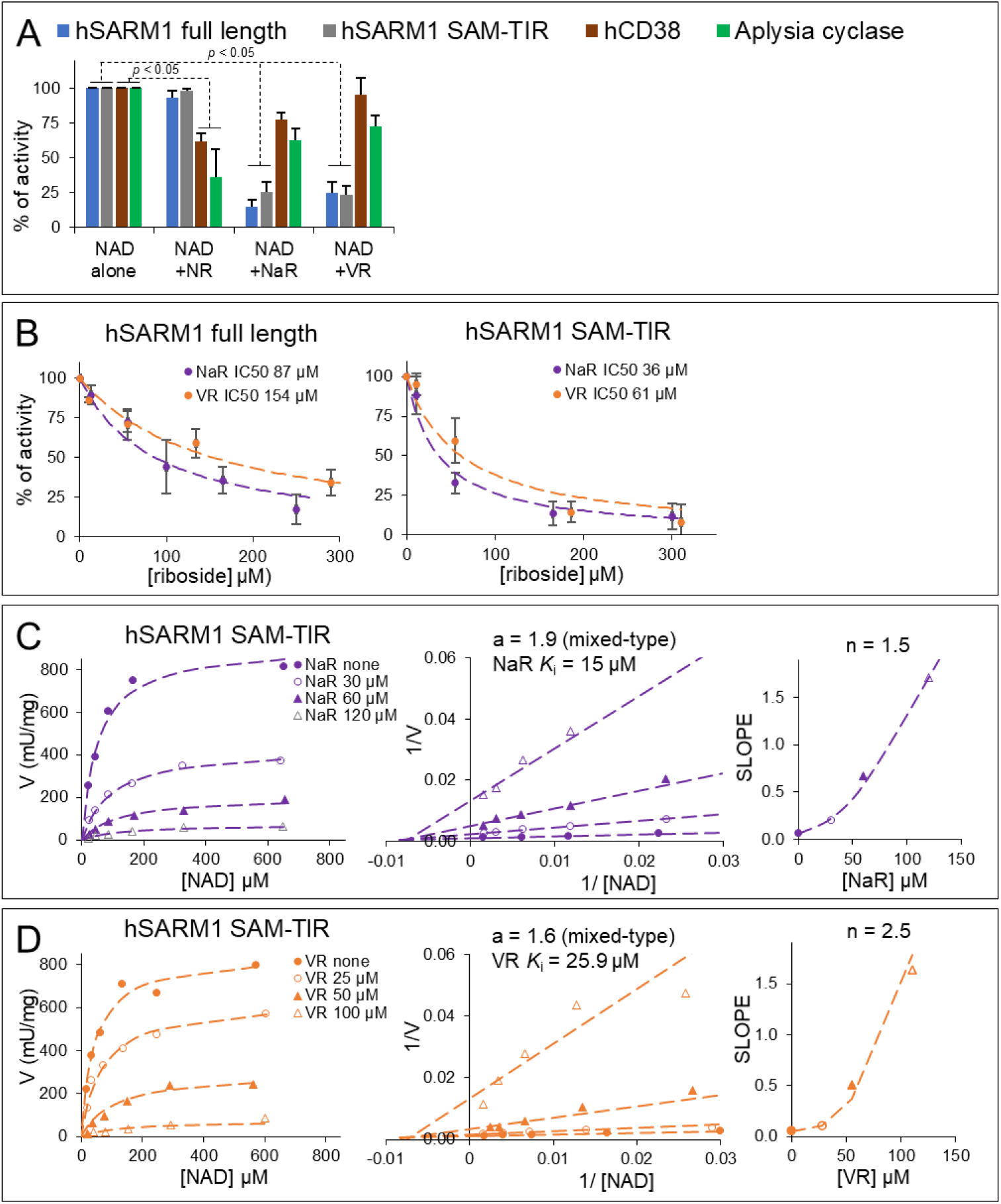
Selective inhibition by pyridine ribosides on multifunctional NAD(P)ase members. (**A**) NADase inhibition at 200 µM fixed NR, NaR, or VR of the indicated enzyme species. Assays were carried out by HPLC at 25 °C for 1 hour using 0.25 mM NAD and 13 µg/ml SARM1 or 0.25 mM NAD and 3.9 µg/ml SAM-TIR or 0.25 mM NAD and 0.1 µg/ml CD38 or 1 mM NAD and 0.02 µg/ml *Aplysia* cyclase. Data are Mean ± SEM from n ≥ 4, normalized to NAD alone controls. (**B**) NADase inhibition at variable NaR or VR of SARM1 full length and SAM-TIR (assayed as in A above). Data are Mean ± SEM from n = 3. Dotted lines, best fitting analyses carried out using equation 4 in Methods with the calculated IC50 values highlighted. (**C**) and (**D**) NADase inhibition kinetics under variable substrate NAD and at various fixed concentrations of either NaR or VR as indicated. SARM1 SAM-TIR (0.6 µg/ml in mix) was assayed by HPLC for 1-3 hours at 25 °C. The graph shows also Lineweaver-Burk plots (**C**,**D middle**) and slope replots (**C**,**D right**) indicating, respectively, inhibition type and *K*_i_ for both NaR and VR and the calculated number (n) of inhibitor molecules that bind to the enzyme. Best fitting analysis was done with equation 5 in Methods.

These data indicate that (1) NaR and VR are good inhibitors of SARM1 catalysis with likely multiple interaction sites and no effects on the ARM-mediated allosteric communication of the protein; and (2) NaR is a potential regulator of SARM1 *in vivo* considering its reported physiological level of 18 µM in wild-type yeast cells [53], and, together with VR, is a promising lead structure for selective drug targeting of SARM1, (3) NaR and NR could be used, together with the alternative substrates above, as a basis for discriminating activity of SARM1 and CD38 in mammalian tissue extracts.

## DISCUSSION

Our data show that base exchange activity of SARM1 can be dominant over hydrolysis and cyclisation of both NAD and NADP and always makes a contribution with each base we tested, and particularly prominent with AcPyr and Vacor (Fig. 4D). We also show that all known SARM1 enzyme activities (NADase, NADPase and base exchange for each dinucleotide) are regulated by physiologically-relevant levels of NMN and NAD at neutral pH, in a manner that requires the N-terminal ARM domain, although below 250 µM NAD SARM1 NADase activity actually falls. Additionally, we describe three new inhibitors of SARM1, two of them present in normal physiology. NADP exerts substrate inhibition at physiologically-relevant levels, and both NaR and VR also inhibit. All three of these compounds are likely to act on the catalytic site in the TIR domain as their effects are independent of the allosteric site in the ARM domain.

The dominance of SARM1 base exchange over hydrolysis and cyclisation depends ultimately on the specific base available but is generally consistent with the greater rate of base exchange previously reported when comparing hydrolysis or cyclisation of an NAD substrate with base exchange involving NADP at two different pH values [16]. It could also help to explain why during the first 4h following axotomy in DRG primary cultures the molar rise in axonal cADPR is 7-8 fold less than the molar fall in NAD, with very little of another alternative product ADPR being detected [28]. Thus, while different cADPR measurements in *Sarm1*^-/-^ and wild-type axons, and the SARM1-dependent increase in cADPR after axotomy [28], clearly show that NAD cyclisation is one relevant physiological and pathological activity of SARM1, it is unlikely to be the only one. Base exchanges from SARM1 in particular are capable of even an almost total contribution to NAD decline under certain conditions.

In addition to confirming a previous report that SARM1 base exchange activity can catalyse the formation of the calcium mobilising agent NaADP, we show that this activity is induced by NMN and occurs at neutral pH not just at pH 4.5. This is important because SARM1 is not known to be a lysosomal enzyme and certainly also localises elsewhere in non-acidic compartments [2, 54]. Thus, our data support a key role of SARM1 in the generation of NaADP, which may explain why *Cd38*^-/-^ do not have lower levels of this metabolite [55, 56] even if CD38 can generate NaADP in specific *in vitro* conditions [57]. Importantly, this also means that the reported absence of any detectable influence of cADPR on axon survival [28] does not exclude a key role for calcium mobilisation as the SARM1-dependent activity driving axon degeneration, consistent with the observed SARM1-dependent rise in calcium [13]. The roles of NaADP and other potential products of base exchange have been relatively neglected by the focus on NAD loss but our findings here show the vital importance of taking a wider perspective of SARM1 activities.

Another SARM1 activity with strong potential to cause axon degeneration is NADP hydrolysis (or cyclisation) [36]. The reduced form of this coenzyme, NADPH, has a crucial role in ROS buffering so SARM1-dependent NADPase puts axons at considerable risk of ROS-induced degeneration, and indeed activation of Wallerian degeneration by vincristine does cause an increase in ROS [58]. Importantly therefore, we now show that SARM1 NADPase as well as NADase and base exchange at neutral pH, is regulated by the NMN:NAD ratio at physiologically-relevant levels of both metabolites. Thus, each, or any combination of these activities could underlie the NMN-dependent degeneration we reported previously [12, 14].

Our data also have many implications for the development of therapies for axonopathies. Animal and cell culture data strongly implicate SARM1-dependent axon loss in some peripheral neuropathies, glaucoma, Parkinson’s disease and other conditions [1, 3, 59] and human genetic studies support its involvement in rare axonopathies involving *NMNAT2* mutation [7, 8] and in ALS [5, 6]. This, together with the complete rescue of axons by SARM1 removal when the pathway is very specifically activated [10, 20] has led to considerable focus on inhibiting and knocking down SARM1 as a therapeutic strategy [3, 28, 32, 60]. Many studies are focusing on blocking the NADase activity specifically but our data now suggest the importance of targeting NADPase and base exchange too. This is important because the most effective drugs will come from targeting the right enzyme activity, indeed small molecules are already known to influence different SARM1 activities in different ways [16, 32].

New physiological inhibitors of SARM1 activity identified in this study indicate potential novel routes to block SARM1. In particular, substrate inhibition by physiologically-relevant levels of NADP suggest that strategies to elevate NADP *in vivo* could be useful. The riboside NaR also has considerable potential on account of its membrane permeability and tolerance by normal physiology, and indeed inhibition of SARM1 may have contributed to its protective role in a reported combinatorial treatment with the NAMPT inhibitor FK866 [61]. However, while NaR is a normal physiological metabolite [53, 62, 63], its levels in most mammalian tissues, including neurons, are uncertain. The deamidated branch of NAD biosynthesis in mammals is a minor component [48] and normal NaR levels are thought to be marginal compared to the historical B3 vitamins Nam and Na and the more recently studied NR [52, 64]. Nonetheless, our *in vitro* finding of a *K*_i_ value in the low micromolar range, lower indeed than *K*_m_ for both substrates (see Fig. 5C,D and Table 1), supports the potential for regulation(s) by NaR *in vivo* under specific conditions, or following exogenous application. Understanding the structural basis of the differential inhibition by NaR, VR, and the inactive NR, will now be a good basis for rational drug design.

Finally, there are several ways in which the preference by SARM1 for base exchange may be exploited as a basis for therapy. This finding suggests that base exchange involving some free bases could contribute substantially to NAD loss, so identifying and lowering the level of such bases could be effective. Our data also indicate a need to reconsider the proposed inhibitory effect of nicotinamide (Nam) on SARM1 NADase [15, 37]. In simple assays of the effect of Nam on NAD hydrolysis it would be impossible to distinguish inhibition from competition between Nam and water in the final step, the former simply switching the incumbent Nam on NAD for a new, identical group and thus appearing to have done nothing. These could only be distinguished using isotopic labelling of substrates. Such an action could also be the basis of the observed therapeutic, or preventative benefit of Nam supply in several conditions involving axon loss [65, 66] so if this could be confirmed as the mechanism there may be ways to boost it further.

In summary, we show that SARM1 is a multidomain NAD(P)ase regulated by pyridine mono and dinucleotides but also by corresponding ribosides and free bases, and that, as shown in Figure 1A, its base exchange may have considerably greater relevance in SARM1-dependent axon death than previously considered. It is regulated like other SARM1 enzyme activities by physiological levels of NMN and NAD, consistent with a role in NMN-dependent axon death [12, 14]; it is likely to be a major physiological source of calcium mobilising signal NaADP; and its manipulation by dietary or pharmacological means has important potential for prevention and therapy of axonal disorders. We also report inhibition of SARM1 by NADP and NaR, with the membrane permeability and physiological tolerance of the latter in particular a property that lends itself to effective therapy.

## SUPPLEMENTAL MATERIAL

**Figure S1.**
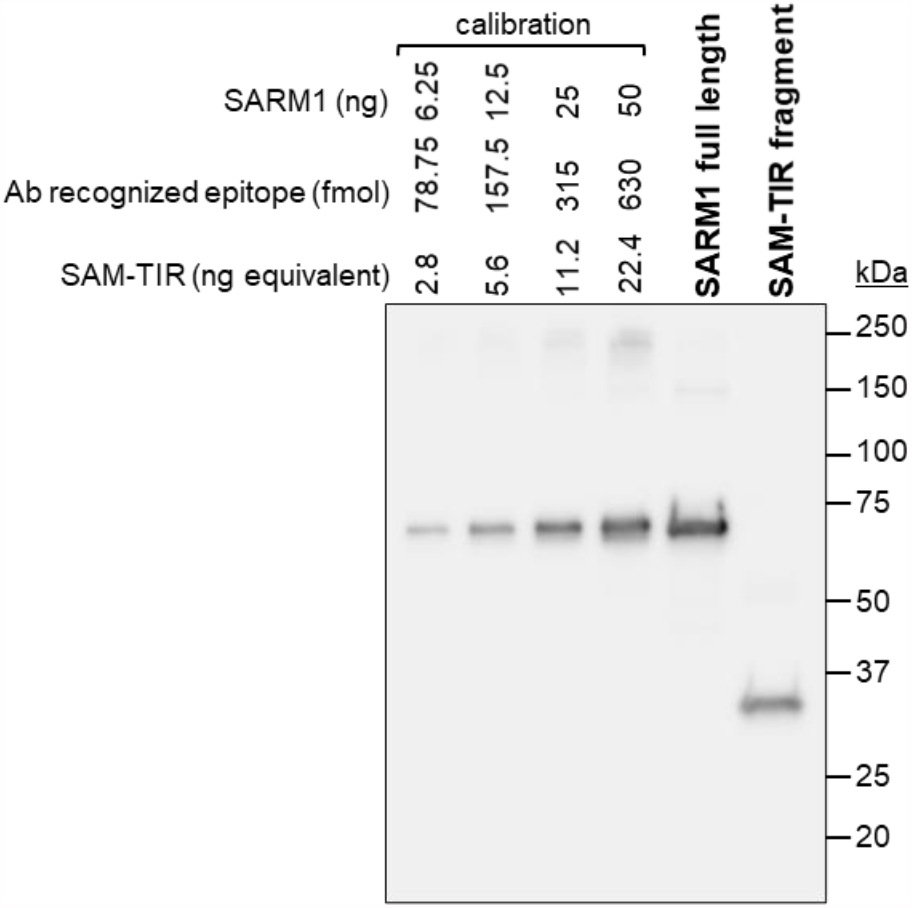
Quantification of human recombinant SARM1 full length and SAM-TIR. Representative immunoblot of SARM1 full length and SAM-TIR (amino acids 409-724) immunoprecipitated from transfected HEK 293T cells. SARM1 and SAM-TIR both possess a C-terminal Flag tag and were immunoprecipitated using a Flag antibody, as described in Methods. The two proteins purified (on beads) were run alongside a dilution series of pure SARM1 standard (not Flag-tagged) to generate a calibration curve to be used for the quantification of both. Then, immunoblots were probed with a polyclonal antibody raised against the SAM-TIR domain of SARM1, namely a protein epitope shared in common by all these proteins. The predicted MWs of 80913.8 (flagged full length) and 37237.2 (flagged SAM-TIR) were used to convert the known amounts of SARM1 (ng) into ng equivalent of SAM-TIR for quantification.

**Figure S2.**
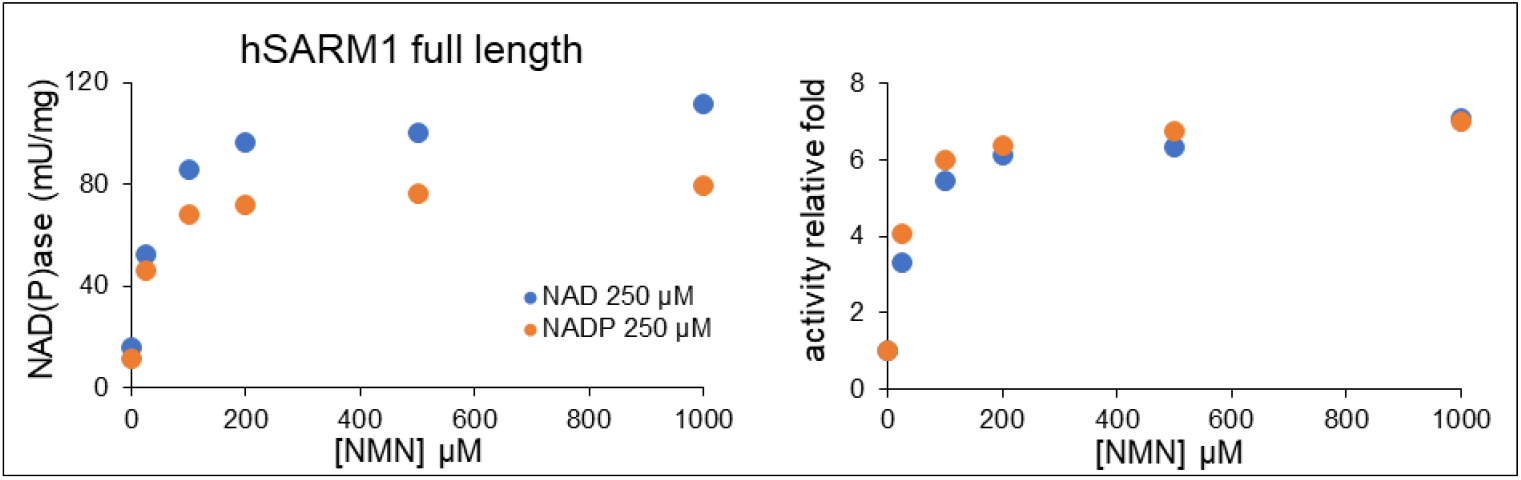
SARM1 NADase and NADPase in parallel after induction by NMN. NADase and NADPase activities of SARM1 full length (14.6 µg/ml per mix) were assayed in comparison by HPLC in the presence of variable NMN as indicated (n = 1). Left graph shows absolute rates while relative ones are on the right. Basal rates without NMN (activity relative fold = 1) were 15.8 mU/mg with 250 µM NAD and 11.3 mU/mg with 250 µM NADP. NMN was not consumed during incubations.

**Figure S3.**
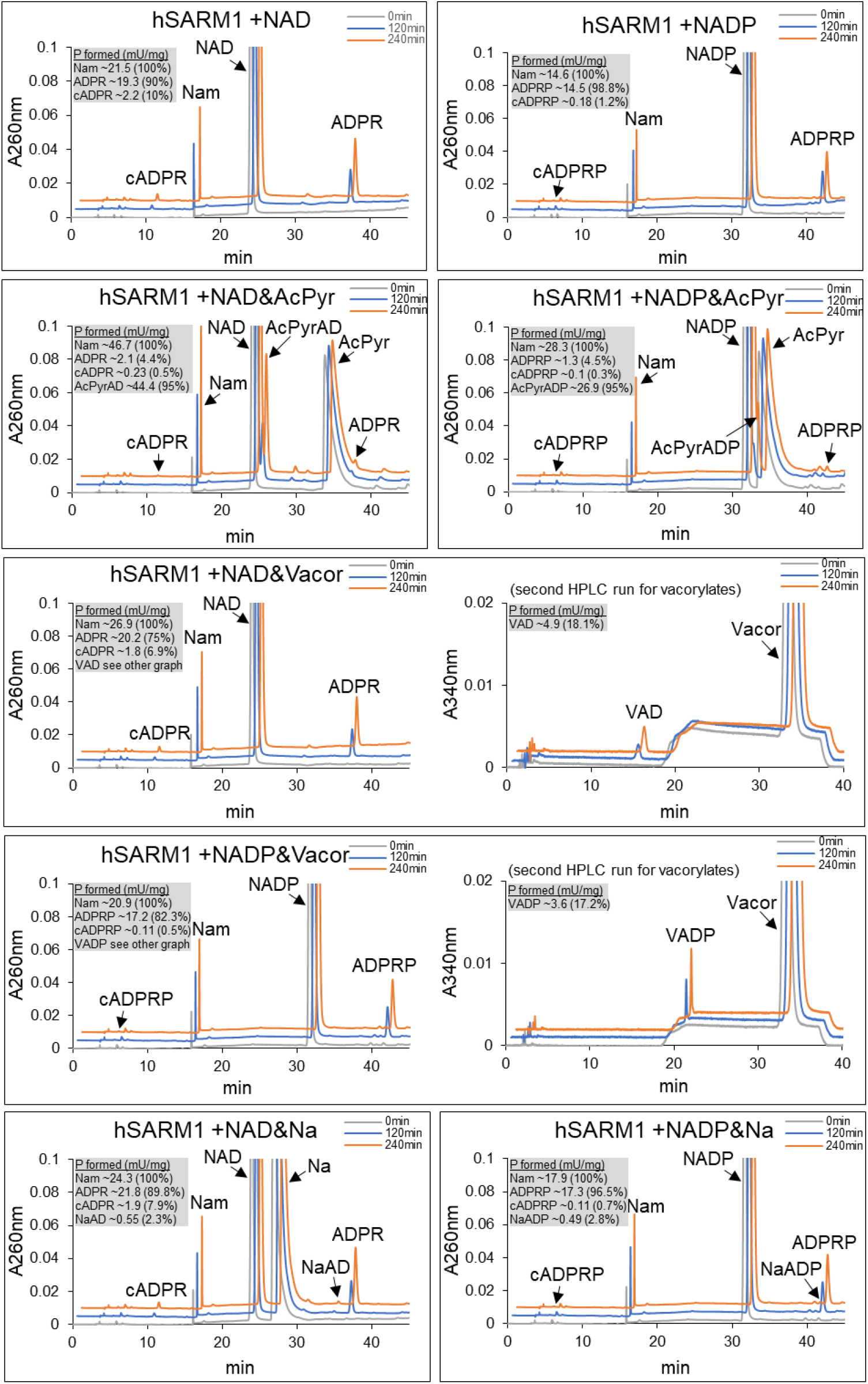
Time course analysis by C18-HPLC of typical base exchange reactions catalysed by SARM1 full length in the presence of single free bases in millimolar excess. Data set of one replicate in Fig. 4B left panel (see assay conditions in the corresponding legend). C18-HPLC analyses were duplicated for vacor compounds (see Methods). Rates in grey boxes are all calculated from the distinct products as indicated, all linearly accumulating by time.

**Figure S4.**
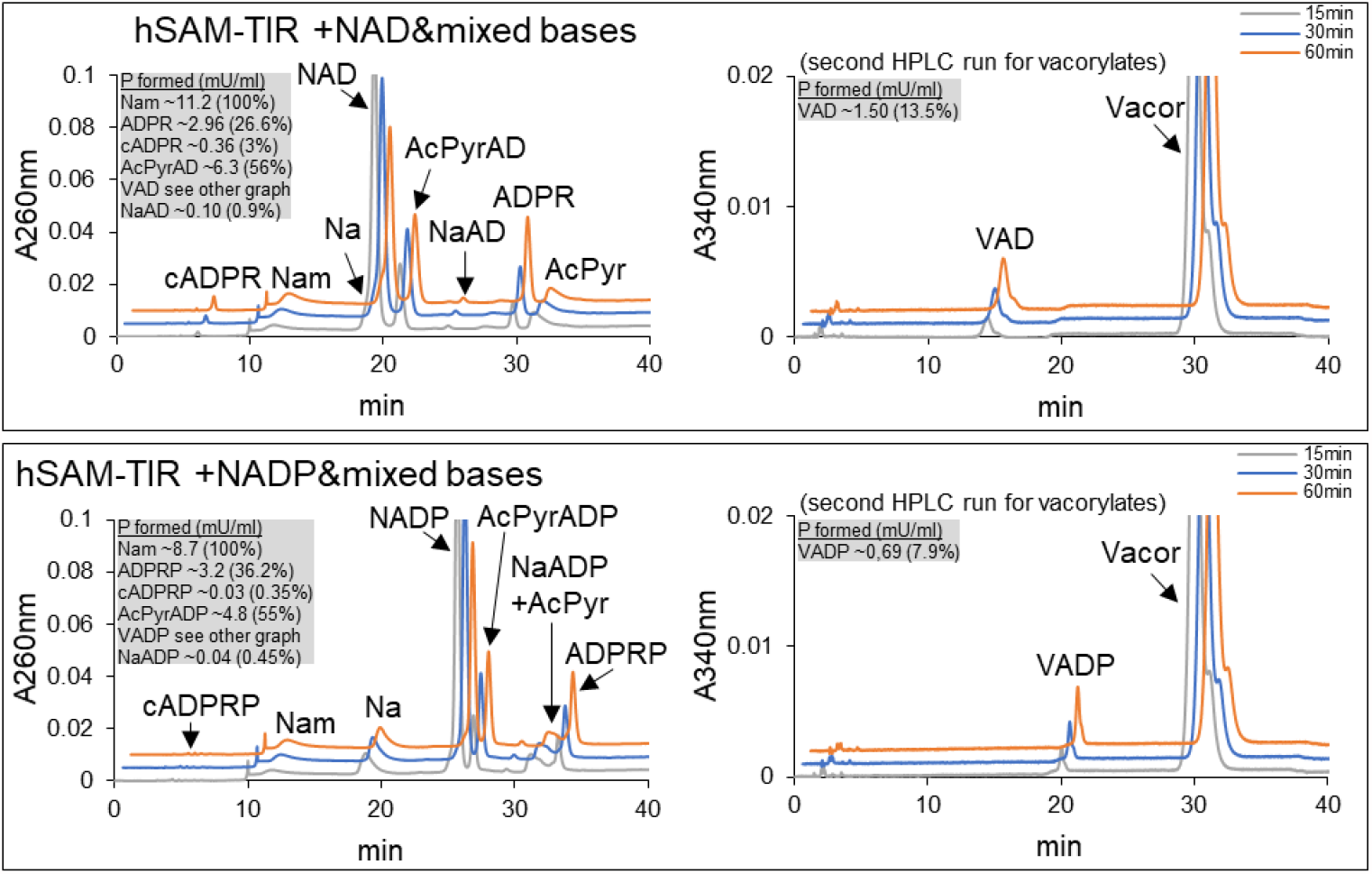
C18-HPLC time course analysis of human SARM1 SAM-TIR base exchange reactions with mixed micromolar concentrations of free bases in the assay mixture. Base exchange reactions at pH 7.5 carried out with ∼5 µg/ml SARM1 SAM-TIR and 250 µM NAD or 250 µM NADP plus free bases mixed together, 3-acetyl pyridine (AcPyr), vacor and nicotinic acid (Na), all three at 250 µM final (equivalent to the substrate). Multiple time stops from both mixtures were collected and analyzed by C18-HPLC showing formation of base exchange products as follows: AcPyrAD from AcPyr and NaAD from Na in the presence of NAD (upper left panel); AcPyrADP from AcPyr and NaADP from Na in the presence of NADP (bottom left panel). The vacor-derived dinucleotide analogs VAD and VADP were measured by duplicated C18-HPLC analysis onto a modified method as described in Methods (right panels). Rates of Nam release indicated in grey boxes corresponded in all cases to the whole substrate consumed, and all other products corresponded to relative percentages of this total as displayed. The assay shows that these three bases lead to formation of corresponding base exchange products even all are present. Furthermore, total base exchange by SARM1 SAM-TIR under these conditions was 70.4% with NAD and of 63.4% with NADP, both in keeping with values obtained when individual bases were used in larger molar excess (see Table S1 and Fig. S3). This reinforces regarding dominance and likely physiological occurrence of SARM1-catalysed base exchange reactions.

**Table S1.**
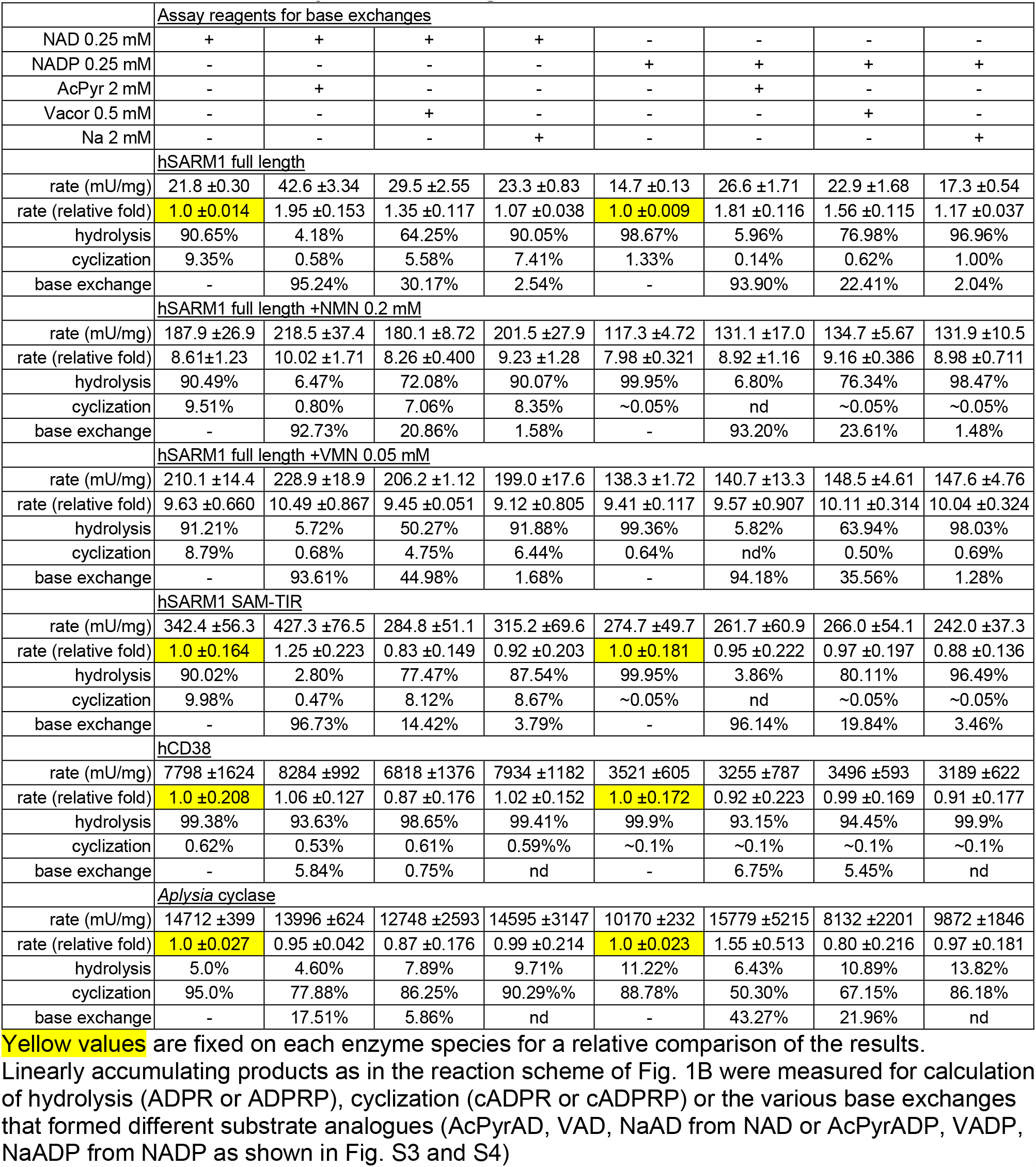
Whole set of data presented in Figure 4.

| Assay reagents for base exchanges |  |  |  |  |  |  |  |  |
| --- | --- | --- | --- | --- | --- | --- | --- | --- |
| NAD 0.25 mM | + | + | + | + | - | - | - | - |
| NADP 0.25 mM | - | - | - | - | + | + | + | + |
| AcPyr 2 mM | - | + | - | - | - | + | - | - |
| Vacor 0.5 mM | - | - | + | - | - | - | + | - |
| Na 2 mM | - | - | - | + | - | - | - | + |
| hSARM1 full length |  |  |  |  |  |  |  |  |
| rate (mU/mg) | 21.8 ±0.30 | 42.6 ±3.34 | 29.5 ±2.55 | 23.3 ±0.83 | 14.7 ±0.13 | 26.6 ±1.71 | 22.9 ±1.68 | 17.3 ±0.54 |
| rate (relative fold) | 1.0 ±0.014 | 1.95 ±0.153 | 1.35 ±0.117 | 1.07 ±0.038 | 1.0 ±0.009 | 1.81 ±0.116 | 1.56 ±0.115 | 1.17 ±0.037 |
| hydrolysis | 90.65% | 4.18% | 64.25% | 90.05% | 98.67% | 5.96% | 76.98% | 96.96% |
| cyclization | 9.35% | 0.58% | 5.58% | 7.41% | 1.33% | 0.14% | 0.62% | 1.00% |
| base exchange | - | 95.24% | 30.17% | 2.54% | - | 93.90% | 22.41% | 2.04% |
| hSARM1 full length +NMN 0.2 mM |  |  |  |  |  |  |  |  |
| rate (mU/mg) | 187.9 ±26.9 | 218.5 ±37.4 | 180.1 ±8.72 | 201.5 ±27.9 | 117.3 ±4.72 | 131.1 ±17.0 | 134.7 ±5.67 | 131.9 ±10.5 |
| rate (relative fold) | 8.61 ±1.23 | 10.02 ±1.71 | 8.26 ±0.400 | 9.23 ±1.28 | 7.98 ±0.321 | 8.92 ±1.16 | 9.16 ±0.386 | 8.98 ±0.711 |
| hydrolysis | 90.49% | 6.47% | 72.08% | 90.07% | 99.95% | 6.80% | 76.34% | 98.47% |
| cyclization | 9.51% | 0.80% | 7.06% | 8.35% | ~0.05% | nd | ~0.05% | ~0.05% |
| base exchange | - | 92.73% | 20.86% | 1.58% | - | 93.20% | 23.61% | 1.48% |
| hSARM1 full length +VMN 0.05 mM |  |  |  |  |  |  |  |  |
| rate (mU/mg) | 210.1 ±14.4 | 228.9 ±18.9 | 206.2 ±1.12 | 199.0 ±17.6 | 138.3 ±1.72 | 140.7 ±13.3 | 148.5 ±4.61 | 147.6 ±4.76 |
| rate (relative fold) | 9.63 ±0.660 | 10.49 ±0.867 | 9.45 ±0.051 | 9.12 ±0.805 | 9.41 ±0.117 | 9.57 ±0.907 | 10.11 ±0.314 | 10.04 ±0.324 |
| hydrolysis | 91.21% | 5.72% | 50.27% | 91.88% | 99.36% | 5.82% | 63.94% | 98.03% |
| cyclization | 8.79% | 0.68% | 4.75% | 6.44% | 0.64% | nd% | 0.50% | 0.69% |
| base exchange | - | 93.61% | 44.98% | 1.68% | - | 94.18% | 35.56% | 1.28% |
| hSARM1 SAM-TIR |  |  |  |  |  |  |  |  |
| rate (mU/mg) | 342.4 ±56.3 | 427.3 ±76.5 | 284.8 ±51.1 | 315.2 ±69.6 | 274.7 ±49.7 | 261.7 ±60.9 | 266.0 ±54.1 | 242.0 ±37.3 |
| rate (relative fold) | 1.0 ±0.164 | 1.25 ±0.223 | 0.83 ±0.149 | 0.92 ±0.203 | 1.0 ±0.181 | 0.95 ±0.222 | 0.97 ±0.197 | 0.88 ±0.136 |
| hydrolysis | 90.02% | 2.80% | 77.47% | 87.54% | 99.95% | 3.86% | 80.11% | 96.49% |
| cyclization | 9.98% | 0.47% | 8.12% | 8.67% | ~0.05% | nd | ~0.05% | ~0.05% |
| base exchange | - | 96.73% | 14.42% | 3.79% | - | 96.14% | 19.84% | 3.46% |
| hCD38 |  |  |  |  |  |  |  |  |
| rate (mU/mg) | 7798 ±1624 | 8284 ±992 | 6818 ±1376 | 7934 ±1182 | 3521 ±605 | 3255 ±787 | 3496 ±593 | 3189 ±622 |
| rate (relative fold) | 1.0 ±0.208 | 1.06 ±0.127 | 0.87 ±0.176 | 1.02 ±0.152 | 1.0 ±0.172 | 0.92 ±0.223 | 0.99 ±0.169 | 0.91 ±0.177 |
| hydrolysis | 99.38% | 93.63% | 98.65% | 99.41% | 99.9% | 93.15% | 94.45% | 99.9% |
| cyclization | 0.62% | 0.53% | 0.61% | 0.59% | ~0.1% | ~0.1% | ~0.1% | ~0.1% |
| base exchange | - | 5.84% | 0.75% | nd | - | 6.75% | 5.45% | nd |
| Aplysia cyclase |  |  |  |  |  |  |  |  |
| rate (mU/mg) | 14712 ±399 | 13996 ±624 | 12748 ±2593 | 14595 ±3147 | 10170 ±232 | 15779 ±5215 | 8132 ±2201 | 9872 ±1846 |
| rate (relative fold) | 1.0 ±0.027 | 0.95 ±0.042 | 0.87 ±0.176 | 0.99 ±0.214 | 1.0 ±0.023 | 1.55 ±0.513 | 0.80 ±0.216 | 0.97 ±0.181 |
| hydrolysis | 5.0% | 4.60% | 7.89% | 9.71% | 11.22% | 6.43% | 10.89% | 13.82% |
| cyclization | 95.0% | 77.88% | 86.25% | 90.29% | 88.78% | 50.30% | 67.15% | 86.18% |
| base exchange | - | 17.51% | 5.86% | nd | - | 43.27% | 21.96% | nd |

**Yellow values** are fixed on each enzyme species for a relative comparison of the results.

## METHODS

### Reagents

Vacor (Pyrinuron N-13738) was purchased from Greyhound Chromatography (UK) and used to generate VMN and VAD enzymatically *in vitro* as reported [20, 67]. The ribosides VR and NaR were also obtained enzymatically from digestion of VMN and NaAD, respectively (see Supplemental Methods). The riboside NR was prepared from Tru Niagen® capsules by dissolving the contents in water and filtering. Other chemicals not indicated otherwise were from Merck, all at the highest purity level and used without further treatment except for NAD, NADP and NaADP that were purified by IEC-FPLC prior to use (see Supplemental Methods).

### Proteins and long-term storage

Human CD38 (P28907, a soluble fragment Val43-Ile300 with 6x-His tag at C terminus) was purchased from R&D Systems (USA). It was diluted to final 0.01 mg/ml in 100 mM HEPES/NaOH buffer, pH 7.5, BSA 0.1 mg/ml and stored at -80 °C in aliquots. *Aplysia californica* ADP ribosyl cyclase was purchased from Merck (A8950), resuspended at 1 mg/ml in Tris/HCl 50mM, pH 7.0 and stored at -20 °C in aliquots. Human recombinant SARM1 full length and the SAM-TIR fragment Val409-Thr724 were expressed as Flag-tagged proteins in HEK 293T cells, immunoprecipitated and quantified by immunoblotting as follows, then stored at -80 °C in aliquots. HEK 293T cells were grown in DMEM with 4,500 mg/L glucose and 110 mg/L sodium pyruvate (PAA), supplemented with 2 mM glutamine, 1% penicillin/streptomycin (both Invitrogen), and 10% fetal bovine serum (PAA). Cells (50-70% confluence) in 10 cm dishes were transfected with 24 µg expression constructs for SARM1 or SAM-TIR [5, 20] using Lipofectamine 2000 reagent (Invitrogen) as per manufacturer instructions. Media was supplemented with 2 mM NR at the time of transfection to boost expression. Transfected cells were collected and washed in ice-cold PBS 24 h after transfection and lysed in ice-cold KHM buffer (110 mM potassium acetate, 20 mM HEPES pH 7.4, 2 mM MgCl2, 0.1 mM digitonin) containing cOmplete™, Mini, EDTA-free protease inhibitor cocktail (Roche) using trituration by pipetting and intermittent vortexing during a 10 min incubation on ice. The use of KHM lysis buffer is critical for subsequent assaying of the true activities of the purified proteins. Lysates were centrifuged for 5 min at 3000 rpm in a microfuge at 4 °C to pellet insoluble material. Supernatants were collected (on ice) and protein concentrations determined using the Pierce BCA assay (Thermo Fisher Scientific). Extracts, diluted to 1 µg/µl in cold KHM buffer, were incubated overnight at 4°C with rotation with 20 µg/ml anti-FLAG M2 antibody (Merck, F3165) and 50 µl/ml of pre-washed Pierce magnetic protein A/G beads (Thermo Fisher Scientific). Beads were collected on a magnetic rack and washed 3x with KHM buffer and 1x with PBS (with protease inhibitors) and then resuspended in PBS containing 1 mg/ml BSA (included to protect proteins during freezing), using ∼150 µl per ml of the immunoprecipitation volume for SARM1 and ∼50 µl for SAM-TIR (due to lower yield resulting from higher enzymatic activity). For immunoblotting, bead suspensions of the immunoprecipitated proteins were diluted 1 : 16 in 2x SDS-PAGE loading buffer and heated to 100 °C for 3 min. Samples (10 µl per well) were resolved on 4-20% gradient gels (Bio-Rad) alongside a dilution series of pure recombinant SARM1 of known concentration (kindly provided by AstraZeneca), before being transferred to Immobilon-P membrane (Millipore). Blots were blocked in TBS (20 mM Tris pH 8.3, 150 mM NaCl) containing 5% skim milk powder for 30 min at RT before being incubated overnight at 4 °C with a human SAM-TIR primary rabbit polyclonal antibody (kindly provided by AstraZeneca) in TBS containing 0.05% Tween-20 (TBST) and 5% milk. After 3x 10 min washes in TBST, blots were incubated for 1-2 h at RT with an anti-rabbit HRP-conjugated secondary antibody (diluted 1 : 3,000, Bio-Rad) in TBST with 5% milk. After 3x 10 min washes in TBST and one rinse in TBS, blots were incubated with Pierce™ ECL Western Blotting Substrate (Thermo Fisher Scientific) and imaged using an Alliance chemiluminescence imaging system (UVITEC Cambridge). Relative band intensities on captured digital images were determined from areas under histogram peaks using Fiji software (http:fiji.sc) and concentrations of SARM1 and SAM-TIR in the bead suspension were calculated from a standard curve generated from the dilution series of a recombinant SARM1 as shown (see Fig. S1).

### NADase assays

hSARM1 was routinely assayed by the NAD/NADH-Glo™ Assay (Promega) while both *Aplysia* cyclase and CD38 were assayed by fluorometry using εNAD [44]. The three NAD(P)ases were also assayed individually in parallel by HPLC in mixtures containing 0.02-11 ug/ml protein and 0.015-4 mM NAD or NADP as substrates, dissolved in 50 mM HEPES/NaOH, pH 7.5. Other reagents used in addition to, or in place of those above are detailed in text or in figure legends, for example metal ions, free bases, dinucleotide analogues, mononucleotides, and ribosides. Assays were all carried out at 25 °C except for temperature-dependence studies. Reactions were stopped by HClO_4_/K_2_CO_3_ and analysed by C18-HPLC as reported [48]. Only the acid labile NADH and NADPH were treated with 1 N NaOH and neutralized using 1:1 vol of 1.73 M KH_2_CO_3_ while the highly hydrophobic metabolites vacor, VAD, VMN and VR, were analysed by a modified C18-HPLC method [67]. Reaction products were quantified from peak areas (see Fig. 2A) based on coelution with standards (except for cyclic products from NAD analogues that were not available and were attributed based on UV-spectra and retention times as predicted). Rates were calculated as follows. Hydrolysis and cyclization were measured from product generation from the respective individual dinucleotide substrate used, e.g., free and cyclic ADP-ribosyl moieties from NAD (Fig. 1B). A basal value for hydrolysis/cyclization was taken as sum of these latter two products. In addition to that value, base exchange or transglycosidation was calculated from the expected dinucleotide analog formed in the presence of the corresponding free base, e.g., AcPyrAD from AcPyr and NAD. The measured level of overall formed products exactly matched in all cases the dinucleotide substrate consumed and the corresponding pyridine base released, according to the reaction scheme (Fig. 1B). Furthermore, enzyme amounts forming 1 µmol/min of overall products above were referred to as single units (U) of activity and initial rates were always considered, for example until a maximum of 20% of the original substrate had been consumed and all products were still accumulating at a nearly linear rate.

Temperature-dependence studies were carried out in buffer as above while pH studies were carried out in universal buffer 50 mM, pH 3-11 (167 mM Tris, 167 mM Bis-Tris, 167 mM sodium acetate). Stocks of 10X universal buffer were adjusted to each pH with 5 N HCl and diluted in the assay. Controls without enzyme were run in parallel to correct for any non-enzymatic NAD degradation at pH ≥ 8.0 or at the respective temperatures studied.

Kinetics were performed by varying NAD or NADP from 15 µM to 4 mM. Their stocks were free from NMN, Nam or any other HPLC-detectable contaminant (see Supplemental Methods), and controls were run in parallel to correct for any potential background. The kinetic parameters were calculated from initial rates obtained from replicates via best-fitting to equations 1 for CD38 and *Aplysia* cyclase, 2 for hSARM1 SAM-TIR, and 3 for hSARM1 full length

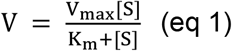

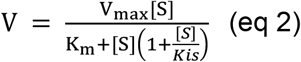

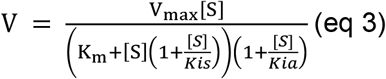

where, for both NAD and NADP, *K*_is_ represents the affinity constant for inhibition on the catalytic TIR site and *K*_ia_ the affinity constant for inhibition on the allosteric ARM site. Kinetics of full length SARM1 with both NAD and NADP was also performed at increasing fixed concentrations of NMN ranging from 5 µM to 50 µM. In that case, the resulting single curves were fit to equation 3 as above to evaluate any competition between NMN and the substrate used (see Table 1).

Inhibition by ribosides was assayed by HPLC initially at fixed 200 µM of each NR, NaR or VR and optimized NAD concentrations for the different enzymes (see Fig. 5 legend). Then, at same NAD concentration as above, IC50 was assessed by varying NaR or VR from 10 µM to 300 µM, and calculated via best fitting of initial rates vs no inhibitor controls (SARM1 activity %) to the equation 4 below. Finally, kinetic constants and mechanism of inhibition were assayed under variable NAD 15-600 µM (where substrate inhibition is negligible) and fixed NaR or VR at various concentrations ranging from 25 µM to 120 µM, and the initial rates obtained were fit to the rate equation 5 below (valid as long as substrate inhibition is negligible)

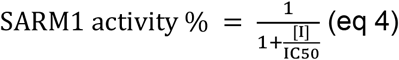

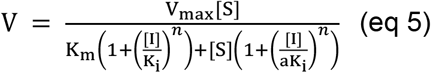

where [I] represents NaR or VR, *K*_i_ is their inhibition constant, “n” is the number of inhibitor molecules that bind the enzyme, “a” is the factor multiplying *K*_i_ that distinguishes the inhibition type, *i*.*e*. competitive (if a = ∞), purely noncompetitive (if a = 1), or mixed (if a<1 or a>1).

Best fitting analyses were done by Excel from the indicated number of replicates and, in most cases, we also presented and confirmed the kinetic parameters obtained from corresponding 1/V vs 1/[S] plots by evaluating the X-and Y-axis intercepts.

### Statistical Analysis

Student’s t test was performed by the Excel software. A *p* value ≤ 0.05 was considered significant and indicated in figure data.

## AUTHOR CONTRIBUTIONS

Conceptualization A.A., G.O.

Methodology A.A., Ca.A., J.G., G.O.

Investigation Ca.A., Ch.A., A.G.T.

Resources & Funding Acquisition M.P.C, G.O.

Writing – Original Draft G.O.

Writing – Review & Editing Ca.A., Ch.A., M.P.C., J.G., A.L.

Supervision & Project development G.O., M.P.C., J.G., A.L.

All authors read and approved the manuscript.

## ACKNOWLEDGMENTS

This work was funded by the BBSRC/AstraZeneca Industrial Partnership Award BB/S009582/1, the Italian Grants RSA 2017-19 from UNIVPM, a Sir Henry Wellcome postdoctoral fellowship from the Wellcome Trust [grant number 210904/Z/18/Z], and the MRC DTP studentship and Gates Foundation.

## COMPETING INTERESTS STATEMENT

Funding for academic research from AstraZeneca and M.P.C. is a consultant for Nura Bio.

## SUPPLEMENTAL METHODS

### Synthesis and purification of Vacor riboside (VR) and Nicotinic acid riboside (NaR)

VR was obtained from the mononucleotide VMN by de-phosphorylation using Alkaline Phosphatase (CIAP). NaR was obtained instead from the dinucleotide NaAD by simultaneous de-phosphorylation/pyrophosphorolysis using both CIAP and Phosphodiesterase I (PDE-I).

### Reagents and reaction mixtures

- **CIAP** (Calf Intestine Alkaline Phosphatase) Applichem A3810 (∼20U/mg at 25 °C) resuspended in 100 mM Tris/HCl, pH 9, 100 mM NaCl to final ∼50mg/ml and stored at -20 °C until use
- **PDE-I** (Phosphodiesterase I from *Crotalus adamanteus*) Merck P3134 (∼0.02U/mg at 37 °C) resuspended in 100 mM Tris/HCl, pH 9, 100 mM NaCl to final ∼10 mg/ml and stored at -20 °C until use
- **VMN** purified as described [20] resuspended in H_2_O to final ∼10 mM (ε_340nm_ of 17.8 mM^-1^ cm^-1^) and stored at -20 °C until use
- **NaAD** Merck N4256 resuspended in H_2_0 to final ∼5 mM and stored at -20 °C until use

|  | Mix (A) for VR |  | Mix (B) for NaR |  |
| --- | --- | --- | --- | --- |
| | ml | final [ ] | μl | final [ ] |
| H<sub>2</sub>O | 202.5 | - | 340 | - |
| Tris/HCl 200mM, pH 9, 200mM NaCl | 250 | 100 mM | 500 | 100 mM |
| MgCl<sub>2</sub> 1 M | 7.5 | 15 mM | 15 | 15 mM |
| CIAP ~50 mg/ml | 20 | ~2 mg/ml | 40 | ~2 mg/ml |
| PDE-I ~10 mg/ml | - | - | 20 | ~0.2 mg/ml |
| VMN ~10 mM | 20(*) | ~0.4 mM | - | - |
| NaAD ~5 mM | - | - | 85(*) | 0.425 mM |
| Final volume | 0.5 ml |  | 1 ml |  |
| Incubation | 1h at 37 °C |  | 2h at 37 °C |  |

Asterisks (\*) indicate last reagents added to start the reactions

Mixtures (A) and (B) after incubation were boiled, centrifuged, and loaded on C18 reverse phase HPLC (Varian, 90 Å, 5 μm). Preparative runs were performed at two distinct temperatures under elution gradients in volatile buffers (see graphs). In particular, VR was eluted at ∼50% acetonitrile after heating the column at 50 °C whereas NaR was eluted at less than 5% acetonitrile after cooling the column at 12 °C. Both ribosides coeluted with corresponding standards (see arrows); they were collected, quantified spectrophotometrically by the ε_340nm_ of 17.8 mM^-1^ cm^-1^ (VR) or the ε_260nm_ of 4.0 mM^-1^ cm^-1^ (NaR), frozen and lyophilized. Dry pellets were stable at -80 °C for months.

### Preparative RP-HPLC of VR and NaR

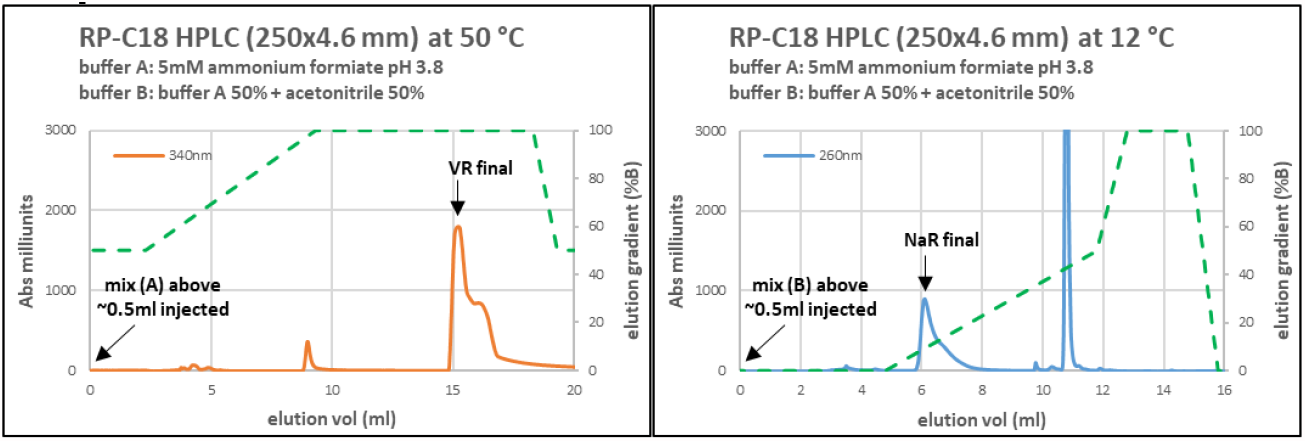

Typical yields by this protocol were ∼150 nmol of VR, *i*.*e*. ∼75% of the original ∼0.2 µmol VMN in mix A, or ∼300 nmol of NaR, *i*.*e*. ∼70% of the original ∼0.43 µmol NaAD in mix B. Lyophilized VR and NaR resulted pure after resuspension in milliQ and re-injection on HPLC in analytical size.

### IEC-FPLC purification of commercial pyridine dinucleotides

Stock solutions in H_2_0 of NAD (Merck N1511), NADP (Merck N5755), and NaADP (Merck N5755) often degraded spontaneously over time or after repeated freeze / thawing, thus generating interfering contaminants such as NMN or Nam. We thus cleaned up these compounds by IEC-FPLC and lyophilized them before using in our SARM1 assays as follows.

For purification, an ion exchange chromatography (IEC) was performed on AKTA Purifier using a TSK DEAE column (Tosoh, 4.6 × 250mm). Equilibration was at 1 ml/min, at 25 °C; elution was under a salt gradient (green dotted lines) as depicted in graphs below. Typically, 1-2 micromoles of NAD or NADP or NaADP per run were injected, collected after elution and mixed from multiple runs, then diluted and quantified by UV absorption (ε_260nm_ of 18 mM^-1^ cm^-1^), frozen and lyophilized. Black arrows in the graphs below indicates most frequent contaminants removed.

### Preparative IEC-FPLC of NAD, NADP, and NaADP and removed contaminants (black arrows)

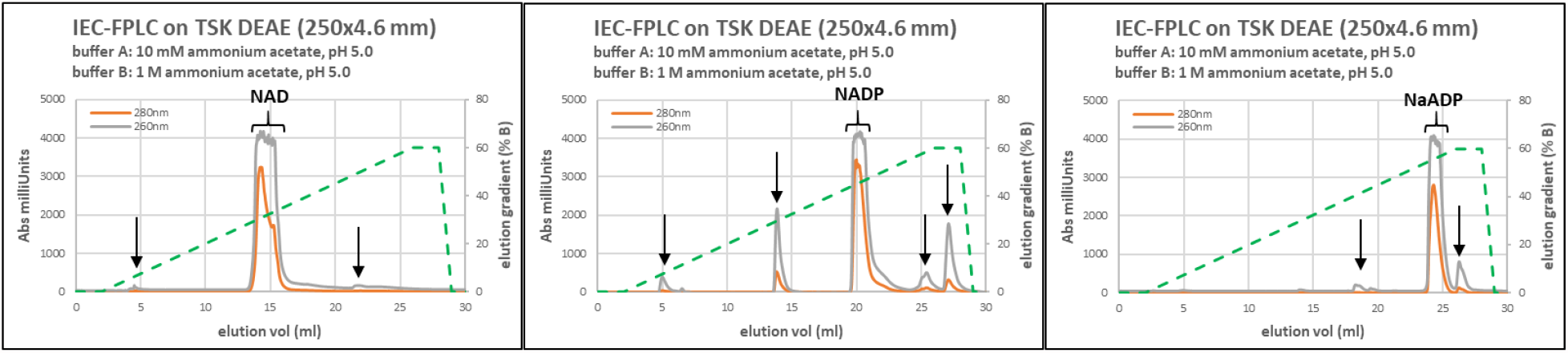

After resuspension and HPLC re-injection, all three dinucleotides above were ≥99.8% pure and were used immediately with no further freezing and thawing.

^1^”Early trials” were defined as the first *N*=2 trials after a novel stimulus was introduced for humans. Relative to the typical interval between insertion trials (*M*≈6 for humans, *M*≈19 for monkeys), this was translated to *N*=6 trials for the monkeys.

#### ABBREVIATIONS

ARM: armadillo/HEAT motif
NAD(P): nicotinamide adenine dinucleotide (phosphate) oxidized forms; this is used throughout to mean
NAD^+^ or NADP^+^; NaAD(P): nicotinic acid adenine dinucleotide (phosphate) oxidized forms; this is used throughout to mean
NaAD^+^ or NaADP^+^; NaR: nicotinic acid riboside
NR: nicotinamide riboside
NAD(P)ase: NAD(P) glycohydrolase (EC 3.2.2.6)
NMNAT2: nicotinamide mononucleotide adenylyltransferase (EC 2.7.7.1) isoform 2
NMN: nicotinamide mononucleotide
ROS: reactive oxygen species
SAM: sterile-α motif
SARM1: sterile alpha and TIR motif-containing protein 1
TIR: Toll/interleukin-1 receptor; vacor or pyrinuron, 1-(4-nitrophenyl)-3-(pyridin-3-ylmethyl)urea
VAD: vacor adenine dinucleotide
VMN: vacor mononucleotide
VR: vacor riboside.

## REFERENCES

1. Coleman, M.P. and A. Hoke, Programmed axon degeneration: from mouse to mechanism to medicine. Nat Rev Neurosci, 2020. 21(4): p. 183–196.

2. Osterloh, J.M., et al., dSarm/Sarm1 is required for activation of an injury-induced axon death pathway. Science, 2012. 337(6093): p. 481–484.

3. Loring, H.S. and P.R. Thompson, Emergence of SARM1 as a Potential Therapeutic Target for Wallerian-type Diseases. Cell Chem Biol, 2020. 27(1): p. 1–13.

4. Conforti, L., J. Gilley, and M.P. Coleman, Wallerian degeneration: an emerging axon death pathway linking injury and disease. Nat Rev Neurosci, 2014. 15(6): p. 394–409.

5. Gilley, J., et al., Enrichment of SARM1 alleles encoding variants with constitutively hyperactive NADase in patients with ALS and other motor nerve disorders. MedRxiv, 2021. https://doi.org/10.1101/2021.06.17.21258268.

6. Bloom, A.J., et al., Constitutively active SARM1 variants found in ALS patients induce neuropathy. BioRxiv, 2021. https://doi.org/10.1101/2021.04.16.439886.

7. Lukacs, M., et al., Severe biallelic loss-of-function mutations in nicotinamide mononucleotide adenylyltransferase 2 (NMNAT2) in two fetuses with fetal akinesia deformation sequence. Exp Neurol, 2019. 320: p. 112961.

8. Huppke, P., et al., Homozygous NMNAT2 mutation in sisters with polyneuropathy and erythromelalgia. Exp Neurol, 2019. 320: p. 112958.

9. Gilley, J., et al., Absence of SARM1 rescues development and survival of NMNAT2-deficient axons. Cell Rep, 2015. 10(12): p. 1974–1981.

10. Gilley, J., R.R. Ribchester, and M.P. Coleman, Sarm1 Deletion, but Not Wld(S), Confers Lifelong Rescue in a Mouse Model of Severe Axonopathy. Cell Rep, 2017. 21(1): p. 10–16.

11. Loreto, A., et al., Mitochondrial impairment activates the Wallerian pathway through depletion of NMNAT2 leading to SARM1-dependent axon degeneration. Neurobiol Dis, 2020. 134: p. 104678.

12. Di Stefano, M., et al., NMN Deamidase Delays Wallerian Degeneration and Rescues Axonal Defects Caused by NMNAT2 Deficiency In Vivo. Curr Biol, 2017. 27(6): p. 784–794.

13. Loreto, A., et al., Wallerian Degeneration Is Executed by an NMN-SARM1-Dependent Late Ca(2+) Influx but Only Modestly Influenced by Mitochondria. Cell Rep, 2015. 13(11): p. 2539–2552.

14. Di Stefano, M., et al., A rise in NAD precursor nicotinamide mononucleotide (NMN) after injury promotes axon degeneration. Cell Death Differ, 2015. 22(5): p. 731–742.

15. Essuman, K., et al., The SARM1 Toll/Interleukin-1 Receptor Domain Possesses Intrinsic NAD(+) Cleavage Activity that Promotes Pathological Axonal Degeneration. Neuron, 2017. 93(6): p. 1334–1343.

16. Zhao, Z.Y., et al., A Cell-Permeant Mimetic of NMN Activates SARM1 to Produce Cyclic ADP-Ribose and Induce Non-apoptotic Cell Death. iScience, 2019. 15: p. 452–466.

17. Figley, M.D., et al., SARM1 is a metabolic sensor activated by an increased NMN/NAD(+) ratio to trigger axon degeneration. Neuron, 2021. 109(7): p. 1118–1136.

18. Sporny, M., et al., Structural basis for SARM1 inhibition and activation under energetic stress. Elife, 2020. 9: p. e62021.

19. Jiang, Y., et al., The NAD(+)-mediated self-inhibition mechanism of pro-neurodegenerative SARM1. Nature, 2020. 588: p. 658–663.

20. Loreto, A., et al., Potent activation of SARM1 by NMN analogue VMN underlies vacor neurotoxicity. BioRxiv, 2020. https://doi.org/10.1101/2020.09.18.304261.

21. Sasaki, Y., et al., Nicotinamide mononucleotide adenylyl transferase-mediated axonal protection requires enzymatic activity but not increased levels of neuronal nicotinamide adenine dinucleotide. J Neurosci, 2009. 29(17): p. 5525–5535.

22. Chini, E.N., CD38 as a regulator of cellular NAD: a novel potential pharmacological target for metabolic conditions. Curr Pharm Des, 2009. 15(1): p. 57–63.

23. Malavasi, F., et al., Evolution and function of the ADP ribosyl cyclase/CD38 gene family in physiology and pathology. Physiol Rev, 2008. 88(3): p. 841–86.

24. Lee, H.C. and Y.J. Zhao, Resolving the topological enigma in Ca(2+) signaling by cyclic ADP-ribose and NAADP. J Biol Chem, 2019. 294(52): p. 19831–19843.

25. Liu, Q., et al., Structural basis for enzymatic evolution from a dedicated ADP-ribosyl cyclase to a multifunctional NAD hydrolase. J Biol Chem, 2009. 284(40): p. 27637–27645.

26. Sauve, A.A., et al., The reaction mechanism for CD38. A single intermediate is responsible for cyclization, hydrolysis, and base-exchange chemistries. Biochemistry, 1998. 37(38): p. 13239–13249.

27. Graeff, R., et al., A single residue at the active site of CD38 determines its NAD cyclizing and hydrolyzing activities. J Biol Chem, 2001. 276(15): p. 12169–12173.

28. Sasaki, Y., et al., cADPR is a gene dosage-sensitive biomarker of SARM1 activity in healthy, compromised, and degenerating axons. Exp Neurol, 2020. 329: p. 113252.

29. Berger, F., M.H. Ramirez-Hernandez, and M. Ziegler, The new life of a centenarian: signalling functions of NAD(P). Trends Biochem Sci, 2004. 29(3): p. 111–118.

30. Gerdts, J., et al., SARM1 activation triggers axon degeneration locally via NAD(+) destruction. Science, 2015. 348(6233): p. 453–457.

31. Aksoy, P., et al., Regulation of intracellular levels of NAD: a novel role for CD38. Biochem Biophys Res Commun, 2006. 345(4): p. 1386–13892.

32. Li, W.H., et al., Permeant fluorescent probes visualize the activation of SARM1 and uncover an anti-neurodegenerative drug candidate. Elife, 2021. 10: p. e67381.

33. Prasad, G.S., et al., Crystal structure of Aplysia ADP ribosyl cyclase, a homologue of the bifunctional ectozyme CD38. Nat Struct Biol, 1996. 3(11): p. 957–964.

34. Liu, Q., et al., Crystal structure of human CD38 extracellular domain. Structure, 2005. 13(9): p. 1331–1339.

35. Horsefield, S., et al., NAD(+) cleavage activity by animal and plant TIR domains in cell death pathways. Science, 2019. 365(6455): p. 793–799.

36. Essuman, K., et al., TIR Domain Proteins Are an Ancient Family of NAD(+)-Consuming Enzymes. Curr Biol, 2018. 28(3): p. 421–430.

37. Bratkowski, M., et al., Structural and Mechanistic Regulation of the Pro-degenerative NAD Hydrolase SARM1. Cell Rep, 2020. 32(5): p. 107999.

38. Yan, T., Y. Feng, and Q. Zhai, Axon degeneration: Mechanisms and implications of a distinct program from cell death. Neurochem Int, 2010. 56(4): p. 529–534.

39. Berthelier, V., et al., Human CD38 is an authentic NAD(P)+ glycohydrolase. Biochem J, 1998. 330(3): p. 1383–1390.

40. Nam, T.S., et al., Interleukin-8 drives CD38 to form NAADP from NADP(+) and NAAD in the endolysosomes to mobilize Ca(2+) and effect cell migration. FASEB J, 2020. 34(9): p. 12565–12576.

41. Loring, H.S., et al., Identification of the first noncompetitive SARM1 inhibitors. Bioorg Med Chem, 2020. 28(18): p. 115644.

42. Orsomando, G., et al., Simultaneous single-sample determination of NMNAT isozyme activities in mouse tissues. PLoS One, 2012. 7(12): p. e53271.

43. Yaku, K., et al., Metabolism and biochemical properties of nicotinamide adenine dinucleotide (NAD) analogs, nicotinamide guanine dinucleotide (NGD) and nicotinamide hypoxanthine dinucleotide (NHD). Sci Rep, 2019. 9(1): p. 13102.

44. Orsomando, G., V. Polzonetti, and P. Natalini, NAD(P)(+)-glycohydrolase from human spleen: a multicatalytic enzyme. Comp Biochem Physiol B Biochem Mol Biol, 2000. 126(1): p. 89–98.

45. Loring, H.S., et al., Initial Kinetic Characterization of Sterile Alpha and Toll/Interleukin Receptor Motif-Containing Protein 1. Biochemistry, 2020. 59(8): p. 933–942.

46. Graeff, R.M., et al., Enzymatic synthesis and characterizations of cyclic GDP-ribose. A procedure for distinguishing enzymes with ADP-ribosyl cyclase activity. J Biol Chem, 1994. 269(48): p. 30260–30267.

47. Hellmich, M.R. and F. Strumwasser, Purification and characterization of a molluscan egg-specific NADase, a second-messenger enzyme. Cell Regul, 1991. 2(3): p. 193–202.

48. Mori, V., et al., Metabolic profiling of alternative NAD biosynthetic routes in mouse tissues. PLoS One, 2014. 9(11): p. e113939.

49. Nakahata, Y., et al., Circadian control of the NAD+ salvage pathway by CLOCK-SIRT1. Science, 2009. 324(5927): p. 654–657.

50. Trammell, S.A. and C. Brenner, Targeted, LCMS-based Metabolomics for Quantitative Measurement of NAD(+) Metabolites. Comput Struct Biotechnol J, 2013. 4: p. e201301012.

51. Belenky, P., et al., Nicotinamide riboside and nicotinic acid riboside salvage in fungi and mammals. Quantitative basis for Urh1 and purine nucleoside phosphorylase function in NAD+ metabolism. J Biol Chem, 2009. 284(1): p. 158–164.

52. Bogan, K.L. and C. Brenner, Nicotinic acid, nicotinamide, and nicotinamide riboside: a molecular evaluation of NAD+ precursor vitamins in human nutrition. Annu Rev Nutr, 2008. 28: p. 115–130.

53. Bogan, K.L., et al., Identification of Isn1 and Sdt1 as glucose-and vitamin-regulated nicotinamide mononucleotide and nicotinic acid mononucleotide [corrected] 5’-nucleotidases responsible for production of nicotinamide riboside and nicotinic acid riboside. J Biol Chem, 2009. 284(50): p. 34861–34869.

54. Gerdts, J., et al., Sarm1-mediated axon degeneration requires both SAM and TIR interactions. J Neurosci, 2013. 33(33): p. 13569–13580.

55. Schmid, F., et al., CD38: a NAADP degrading enzyme. FEBS Lett, 2011. 585(22): p. 3544–3548.

56. Soares, S., et al., NAADP as a second messenger: neither CD38 nor base-exchange reaction are necessary for in vivo generation of NAADP in myometrial cells. Am J Physiol Cell Physiol, 2007. 292(1): p. C227–239.

57. Guse, A.H. and H.C. Lee, NAADP: a universal Ca2+ trigger. Sci Signal, 2008. 1(44): p. re10.

58. Press, C. and J. Milbrandt, Nmnat delays axonal degeneration caused by mitochondrial and oxidative stress. J Neurosci, 2008. 28(19): p. 4861–4871.

59. Krauss, R., et al., Axons Matter: The Promise of Treating Neurodegenerative Disorders by Targeting SARM1-Mediated Axonal Degeneration. Trends Pharmacol Sci, 2020. 41(4): p. 281–293.

60. Gould, S.A., et al., Protection against oxaliplatin-induced mechanical and thermal hypersensitivity in Sarm1(-/-) mice. Exp Neurol, 2021. 338: p. 113607.

61. Liu, H.W., et al., Pharmacological bypass of NAD(+) salvage pathway protects neurons from chemotherapy-induced degeneration. Proc Natl Acad Sci U S A, 2018. 115(42): p. 10654–10659.

62. Kulikova, V., et al., Generation, Release, and Uptake of the NAD Precursor Nicotinic Acid Riboside by Human Cells. J Biol Chem, 2015. 290(45): p. 27124–27137.

63. Carpi, F.M., et al., Simultaneous quantification of nicotinamide mononucleotide and related pyridine compounds in mouse tissues by UHPLC-MS/MS. Separation Science Plus, 2018. 1(1): p. 22–30.

64. Bieganowski, P. and C. Brenner, Discoveries of nicotinamide riboside as a nutrient and conserved NRK genes establish a Preiss-Handler independent route to NAD+ in fungi and humans. Cell, 2004. 117(4): p. 495–502.

65. Hui, F., et al., Improvement in inner retinal function in glaucoma with nicotinamide (vitamin B3) supplementation: A crossover randomized clinical trial. Clin Exp Ophthalmol, 2020. 48(7): p. 903–914.

66. Anderson, G.D., et al., The effect of nicotinamide on gene expression in a traumatic brain injury model. Front Neurosci, 2013. 7: p. 21.

67. Buonvicino, D., et al., Identification of the Nicotinamide Salvage Pathway as a New Toxification Route for Antimetabolites. Cell Chem Biol, 2018. 25(4): p. 471–482.

